# PiVR: an affordable and versatile closed-loop platform to study unrestrained sensorimotor behavior

**DOI:** 10.1101/2019.12.20.885442

**Authors:** David Tadres, Matthieu Louis

## Abstract

Tools enabling closed-loop experiments are crucial to delineate causal relationships between the activity of genetically-labeled neurons and specific behaviors. We developed the Raspberry <u>Pi</u> <u>V</u>irtual <u>R</u>eality system (PiVR) to conduct closed-loop optogenetic stimulation of neural functions in unrestrained animals. PiVR is an experimental platform that operates at high-temporal resolution (>50 Hz) with low latencies (~10 ms), while being affordable (<$500) and easy to build (<6 hours). This tool was designed to be accessible to a wide public, from highschool students to professional researchers studying systems neuroscience. We illustrate the functionality of PiVR by focusing on sensory navigation in response to gradients of chemicals (chemotaxis) and light (phototaxis). We show how *Drosophila* flies perform negative chemotaxis by modulating their locomotor speed to avoid locations associated with optogenetically-evoked bitter taste. In *Drosophila* larvae, we use innate positive chemotaxis to compare orientation behavior elicited by real- and virtual-odor gradients with static shapes as well as by turbulent virtual-odor plumes. Finally, we examine how positive phototaxis emerges in zebrafish larvae from the modulation of turning maneuvers during the ascent of virtual white-light gradients. Besides its application to study chemotaxis and phototaxis, PiVR is a versatile tool designed to bolster efforts to map and to functionally characterize neural circuits.

## Introduction

Behavior emerges from the conversion of sensory input into motor output. This process, called sensorimotor transformation (1), results from the integrated activity of neural circuits in the nervous system (2). Since the advent of molecular tools to map and manipulate the activity of genetically-targeted neurons (3, 4), a major goal of systems neuroscience has been to crack the neural computations underlying sensorimotor transformation (5, 6). Due to the probabilistic nature of behavior (7–9), probing sensorimotor functions requires stimulating an animal repeatedly with reproducible patterns of sensory input. These conditions can be achieved by immersing animals in virtual realities (10).

A virtual-reality paradigm consists of a simulated environment perceived by an animal and updated based on a readout of its behavior. Historically, virtual realities have been introduced to study optomotor behavior in tethered flies and bees (11, 12). For several decades, sophisticated computer-controlled methods have been developed to produce ever more realistic immersive environments. In FreemoVR, freely-moving flies avoid collisions with fictive tridimensional obstacles and zebrafish engage in social interactions with artificial peers (13). Spatial learning has been studied in tethered flies moving on a treadmill in two-dimensional environments filled with geometrical objects projected on a visual display (14). The same technology has been used to record the neural activity of mice exploring a virtual space (15). Although virtual realities were initially engineered to study visual behavior, they have been generalized to other sensory modalities such as touch and olfaction. In a treadmill system, navigation has been studied in mice directed by localized stimulations of their whiskers (16). The combination of closed-loop tracking and optogenetic stimulation of genetically-targeted sensory neurons has enabled a quantitative analysis of chemotaxis in freely-moving *C. elegans* and in *Drosophila* larvae immersed in gradients (17, 18).

Virtual-reality assays aim to reproduce the natural feedback that binds behavior to sensation (10). First, the behavior of an animal must be accurately classified in real time. In tethered flying flies, wing beat patterns have been used to deduce turning maneuvers (19). Likewise, the movement of a tethered walking fly or a mouse can be inferred from the rotation of the spherical trackball of a treadmill (15, 20). In freely moving *C. elegans* and *Drosophila* larvae, the posture of an animal and the position of specific body parts —the head and tail, for instance-have been tracked during motion in two-dimensional arenas (17, 18). Variables related to the behavioral readout —most commonly, the spatial coordinates— are then mapped onto a virtual sensory landscape to update the stimulus intensity (15). The effectiveness of the virtual-reality paradigm is conditioned by the overall temporal delay between the animal’s behavior and the update of the stimulus. The shorter this delay, the more authentic the virtual reality is perceived.

Irrespective to the model organism under study, the methodology deployed to create efficient virtual realities with closed-loop tracking relies on advanced technology that makes behavioral setups costly and often difficult to adopt by non-specialists. There is a pressing need to complement the existing collection of sophisticated assays with tools that are affordable and accessible to most labs. The fly “ethoscope” proposes a hardware solution exploiting 3D printing to study the behavior of adult flies (21). In this system, tracking and behavioral classification are implemented by a portable and low-cost computer, the Raspberry Pi. Although this system can implement a feedback loop between real-time behavioral tracking and stimulus delivery (e.g., the physical rotation of an assay to disrupt sleep), it was not conceived to create refined virtual-reality environments using optogenetic stimulations.

Here, we present PiVR, a Raspberry <u>Pi</u> <u>V</u>irtual <u>R</u>eality platform enabling the presentation of virtual realities to freely-moving small animals. This closed-loop tracker was designed to be accessible to a wide range of researchers by keeping the construction costs low and by maintaining the basic operations simple and customizable to suit the specificities of new experiments. We benchmark the performance of PiVR by studying navigation behavior in the *Drosophila* larva. We then reveal how adult flies adapt their speed of locomotion to avoid areas associated with the activation of bitter sensing neurons. Next, we show that zebrafish larvae approach a virtual-light source by modifying their turn angle in response to temporal changes in light intensity. Finally, we establish that PiVR can create dynamic sensory landscapes to study navigation in turbulent odor plumes.

## Results

### PiVR permits high-performance closed-loop tracking and optogenetic stimulations

The PiVR system enables high-resolution, optogenetic, closed-loop experiments with small freely-moving animals. In its standard configuration, PiVR is composed of a behavioral arena, a camera, a Raspberry Pi microcomputer, a LED controller, and a touchscreen (Figure 1A). The platform is controlled via a user-friendly graphical interface (Figure 1B). The material for one setup amounts to less than $500 with the construction costs decreasing to about $350 when several units are built in parallel (S1 Table). In spite of its affordability, PiVR runs a sophisticated customized software (Supplementary Figures S1-6) that automatically identifies semitransparent animals such as *Drosophila* larvae embedded in a static background (Figure 2) and that monitors movement at a frame rate sufficient to accurately track rapidly-moving animals such as walking adult flies and zebrafish larvae (Figures 3 and 4).

**Figure 1:**
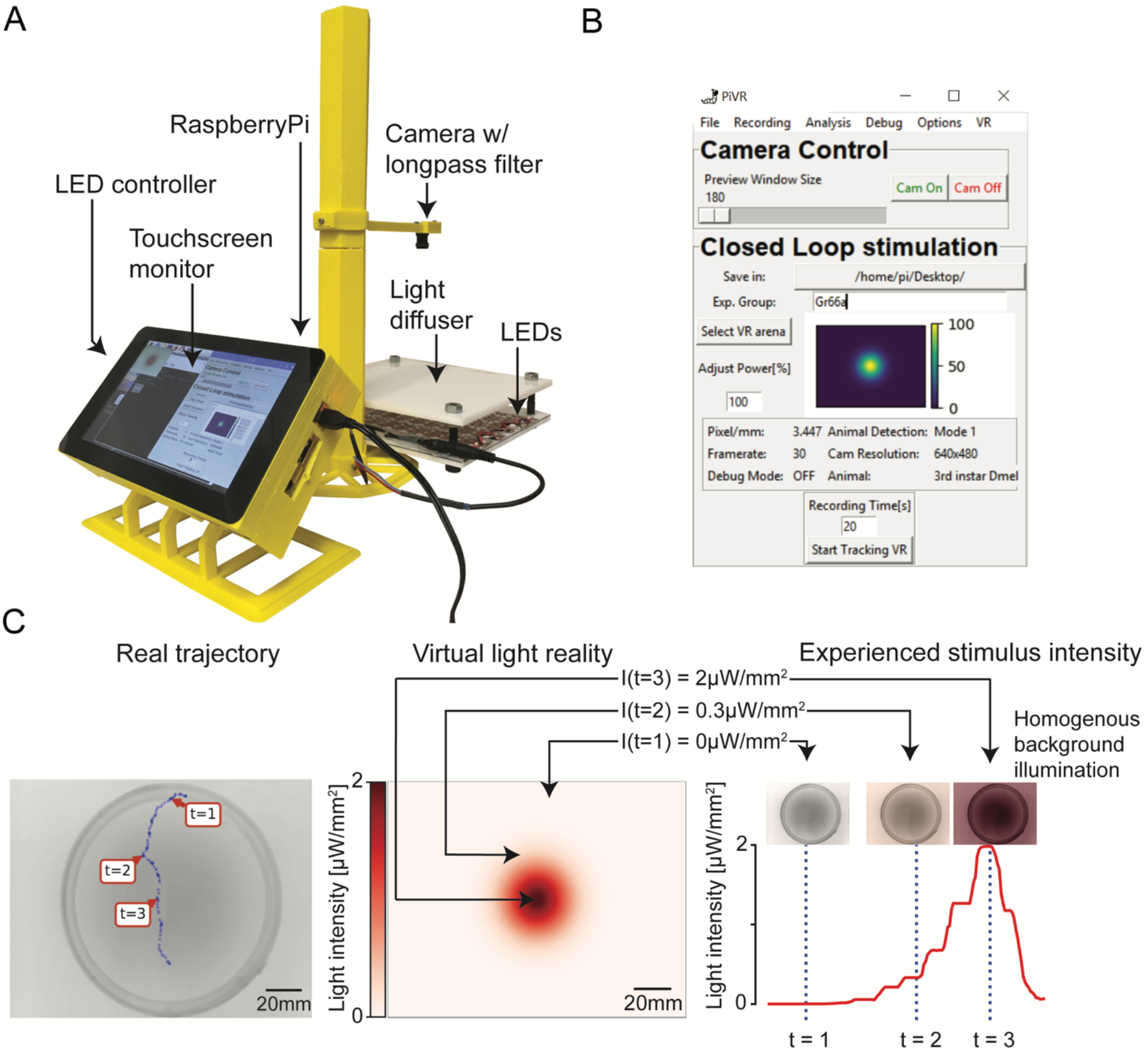
Virtual realities created by PiVR. **(A)** Picture of the standard PiVR setup. The animal is placed on the light diffuser and illuminated from below using infrared LEDs and recorded from above. The Raspberry Pi computer and the LED controller are attached to the touchscreen, which permits the user to interface with the PiVR setup. **(B)** Screenshot of the graphical user interface (GUI) while running a virtual reality experiment. The GUI has been designed to be intuitive and easy to use while presenting all important experimental parameters that can be modified. **(C)** Virtual realities are created by updating the intensity of a homogeneous light background based on the current position of a tracked animal mapped onto predefined landscape shown at the center. (Center) Predefined virtual gradient with a Gaussian geometry. (Left) Trajectory of an unconstrained animal moving in the physical arena. (Right) The graph indicates the time course of the light intensity experienced by the animal during the trajectory displayed in the left panel. Depending on the position of the animal in the virtual-light gradient, the LEDs are turned off (t=1), turned on at an intermediate (t=2) or maximum intensity (t=3).

PiVR has been designed to create virtual realities by updating the intensity of a homogeneous stimulation backlight based on the current position of a tracked animal (Figure 1C, left panel) relative to a preset landscape (Figure 1C, middle panel, S1 Movie). Virtual sensory realities are generated by optogenetically activating sensory neurons of the peripheral nervous system (3). In the present study, we focus on applications involving CsChrimson since the red activation spectrum of this light-gated ion channel is largely invisible to *Drosophila* (22). Depending on the light-intensity range necessary to stimulate specific neurons, PiVR features a light pad emitting stimulation light at low-to-medium (2 μW/mm^2^) or high (22 μW/mm^2^) intensities (Supplementary Figures S2C and S2D). A key advantage of the closed-loop methodology of PiVR is that it permits the creation of virtual realities with arbitrary properties free of the physical constraints of real stimuli. In Figure 2Fi, we illustrate the use of PiVR by immersing *Drosophila* larvae in a virtual-odor gradient that has the shape of a “volcano”.

**Figure 2:**
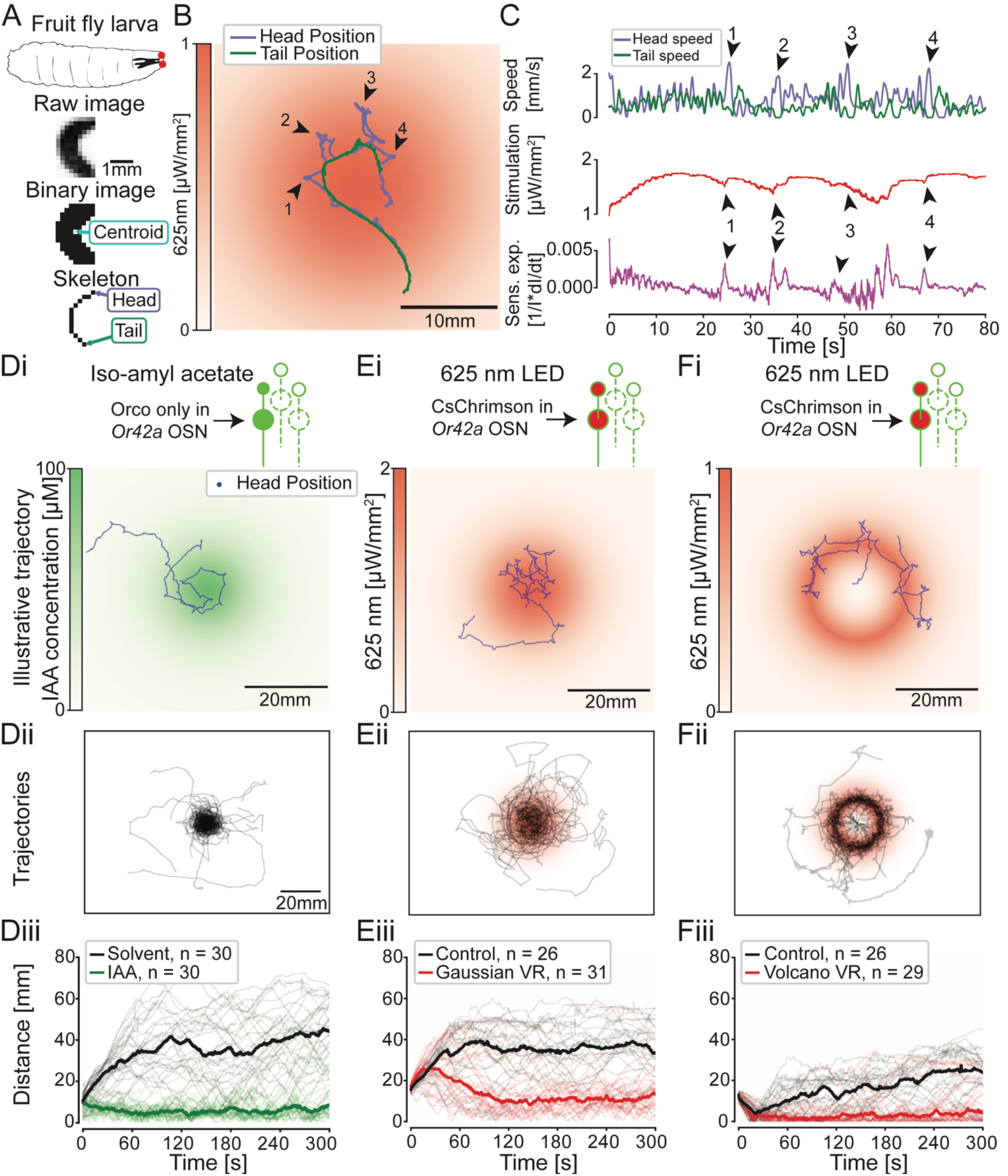
Benchmarking PiVR performance by eliciting larval chemotaxis in virtual-odor gradients. **(A)** *Drosophila* larva with a pair of single *Or42a*-functional olfactory sensory neurons (OSNs) (red dots). Illustration of the identification of different body parts of a moving larva by PiVR. (B) Illustrative trajectory of a larva in a Gaussian virtual-odor gradient elicited by light stimulation. Arrowheads and numbers indicate lateral head movements (casts) and the timepoints are congruent with the arrowheads shown in (C). Panel (D) shows the beh avior of *Drosophila* larvae directed by *Or42a* OSNs in a gradient of isoamyl acetate (green color). (E-F) Behavior of larvae expressing the light-gated ion channel CsChrimson in the *Or42a* olfactory sensory neurons evoking a virtual-odor gradient. In panel (E), the virtual-odor gradient (red) has a geometry similar to the real-odor gradient (green) presented in (D). The “volcano” virtual-odor landscape presented in (F) highlights that the information conveyed by the *Or42a* OSN alone is sufficient for larvae to chemotax with high accuracy along the rim of the gradient. Thick lines in panels Diii, Eiii and Fiii indicate the median distances to source and the light traces indicate individual trials.

The ability to create virtual sensory experiences that appear authentic to a test subject critically depends on the time (latency) that elapses between an action of the subject, its detection by the tracking software and the update of the light intensity. Latencies that are too long can disrupt natural sensorimotor feedback leading to confusion (23). We characterized the overall latency of PiVR by measuring the following three parameters: (i) image acquisition time, (ii) image processing time and (iii) the time taken for commands issued by the Raspberry Pi to be actuated by the LED hardware (Supplementary Figure S1). We find that the image processing time is the main time-consuming step. The total latency between the moment an image starts being recorded and the intensity of the LED system is updated based on the analysis of that image is shorter than 20 ms (Supplementary Figure S1Cii and S1Dii). Thus, PiVR is suited to perform on-line tracking of small animals at a frame rate of 50 Hz, which has been shown to be sufficiently short to create virtual olfactory realities in *Drosophila* larvae (17) and virtual visual realities in walking adult flies (20).

The PiVR software implements image acquisition, object tracking, and the update of background illumination for optogenetic stimulation. It is free, fully open-source and written in the programming language Python. At the beginning of each experiment, an auto-detect algorithm separates the moving object —the animal— from the background (Supplementary Figure S3). During the rest of the experiment, the tracking algorithm operates based on a principle of local background subtraction to achieve high frame rates (Supplementary Figure S4). Besides locating the position of the animal’s centroid, PiVR uses a Hungarian algorithm to tell apart the head from the tail positions (Supplementary Figure S5). For applications involving off-line tracking with a separate software (24, 25), the on-line tracking module of PiVR can be disabled to record videos at 90 frames per second in an open-loop mode.

The possibility to 3D printing components that once required sophisticated machining has empowered the “maker movement” in our scientific community (26). Inspired by this philosophy, PiVR is built from hardware parts that are 3D printed. Thus, the modular design of the setup can be readily adapted to accommodate the experimental needs of traditional model organisms (larvae, adult flies and zebrafish) as well as less conventional small animals (Supplementary Figures S2 and S6). For example, we adapted PiVR to acquire movies of 10 fruitfly larvae simultaneously with an image quality sufficient to permit off-line tracking of multiple animal tracking with the idtracker.ai software (25). To achieve this, we modified the design of the arena to allow illumination from the side instead of bottom (Supplementary Figure S2B). This adaptation of the illumination setup was necessary to enhance contrast in the appearance of individual larvae for idtracker.ai to detect idiosyncratic differences between larvae (Supplementary Figure S2Bi). The versatility of PiVR was also illustrated by tracking various arthropods and a vertebrate with diverse body structures and locomotor properties (Supplementary Figure S6, Figures 2, 3, 4 and 5).

**Figure 3:**
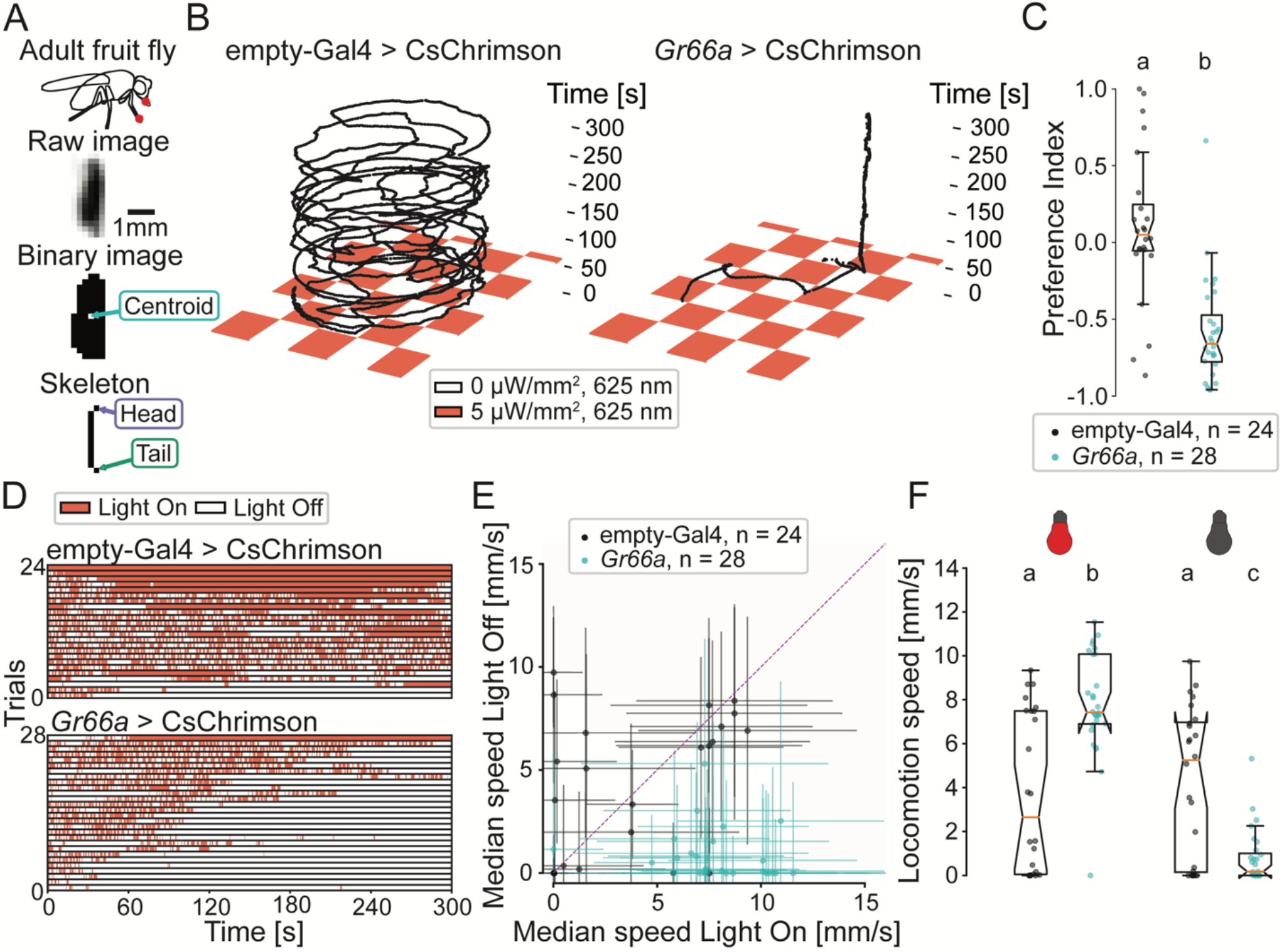
Adult fruit flies avoid activation of bitter-sensing neurons by modulating their locomotion speed. **(A)** Adult *Drosophila* expressing CsChrimson in *Gr66a* bitter sensing neurons (red circles). Illustration of the identification of different body parts of a moving fly by PiVR. **(B)** Illustrative trajectories of flies in a virtual checkerboard pattern and the corresponding ethogram. Flies were behaving in a petri dish. **(D)**. The ethogram reports the time spent by individual animals (rows) in the dark (white) and lit (red) squares. Panel **(C)** displays a quantification of the avoidance of virtual bitter taste through a preference index: 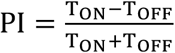 where T is the time spent on the ON or OFF quadrants (Mann-Whitney U Test, p < 0.001). **(E)** Median locomotion speeds of individual animals as a function of the exposure to light. **(F)** Quantification of locomotion speeds across experimental conditions (Dunn’s multiple comparisons test, different letters indicate at least p < 0.01). Statistical procedures are detailed in the Methodology section. Statistical significances are indicated with lowercase letters.

**Figure 4:**
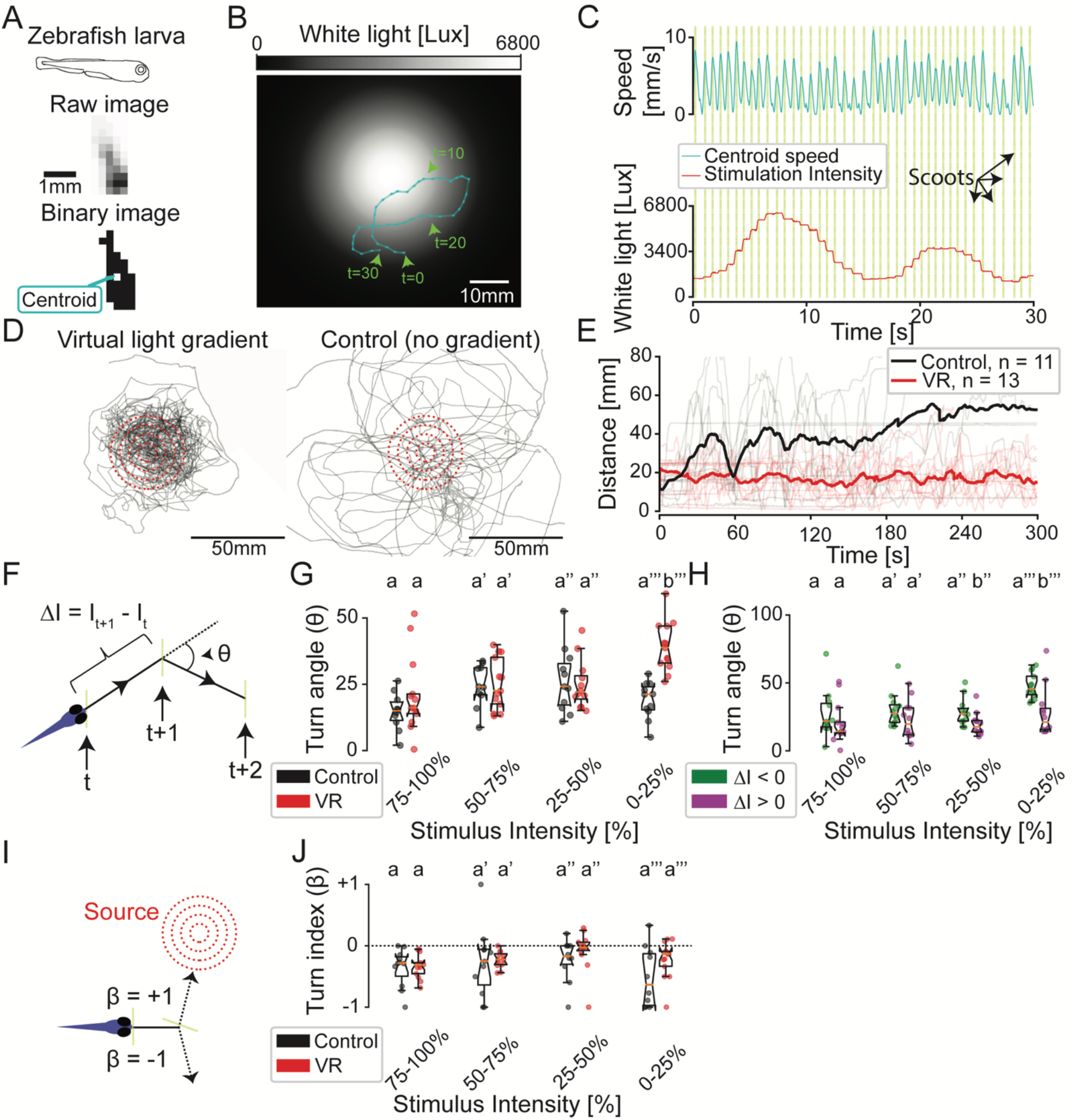
Zebrafish larvae adapt their turn dynamics to stay close to a virtual-light source. **(A)** Illustration of the identification of a moving zebrafish larva by PiVR. **(B)** Illustrative trajectory of a zebrafish larva in a virtual-light gradient having a Gaussian geometry. Panel **(C)** displays the time course of the speed and the white-light intensity during that trajectory shown in panel **(B)**. Yellow vertical lines indicate automatically detected scoots. **(D)** Trajectories of 13 fish tested in virtual-light gradient **(left)** and 11 fish in control **(right).** Red circles indicate 10, 20, 30 and 40 mm distance to the center to the virtual-light source (see Methodology). **(E)** Thick lines indicate the time courses of the median distances to the virtual-light source. The light lines indicate individual trials. **(F)** Illustration of the discretization of a trajectory segment into scoots: yellow vertical lines indicate the position where the animal stops, reorients and starts the next scoot. Black dashed lines indicate movement of the fish. The turn angle (θ) and change in light intensity (ΔI) are calculated for every pair of consecutive scoots (see Methodology). **(G)** Relationship between θ and ΔI during the previous scoot (Independent two-sample t-test, different letters indicate p < 0.001). **(H)** Turn angles θ of the virtual reality condition are grouped according to negative (green) and positive (magenta) intensity experienced in the previous scoot (t-test for paired samples, different letters indicate at least p < 0.05). **(I)** The turn index (β) is calculated from the orientation of the animal relative to the virtual-light source. **(J)** Turn index (β) as a function of stimulus intensity (Mann-Whitney U Test, all groups p > 0.05). All reported statistical significances are Bonferroni corrected and indicated with lowercase letters. Statistical procedures are detailed in the Methodology section.

### Benchmarking PiVR performances by eliciting larval chemotaxis in virtual-odor gradients

To demonstrate the use of PiVR, we turned to the navigation behavior evoked by airborne-odor gradients (chemotaxis) in the *Drosophila* larva (27, 28). Larval chemotaxis relies on a set of well-characterized sensorimotor rules (29). To ascend an attractive odor gradient, larvae modulate the alternation of relatively straight runs and reorientation maneuvers (turns). Stops are predominantly triggered when the larva undergoes negative changes in odor concentration during down-gradient runs. Following a stop, turning is directed towards the gradient through an active sampling process that involves lateral head movements (head casts). A key advantage of the larva as a model organism for chemotaxis is that robust orientation responses can be directed by a functionally reduced olfactory system. The *Or42a* odor receptor gene is expressed in a pair of bilaterally symmetric olfactory sensory neurons (OSNs) (30, 31) that are sufficient to direct chemotaxis (32). We exploited this property to compare reorientation performances elicited by real and virtual-odor stimulations of the *Or42a* OSNs.

We started by applying the computer-vision algorithm of PiVR to track larvae with a single functional *Or42a*-expressing OSN (Figure 2A). Individual animals were introduced in a rectangular arena comprising a gradient of isoamyl acetate (IAA) at its center (Figure 2B). Once exposed to the odor gradient, *Or42a*-functional larvae quickly identified the position of the odor source and they remained in the source’s vicinity (Figures 2B, 2D and S2 Movie). The trajectories of consecutively-tested larvae were analyzed by quantifying the time course of the distance between the head of the larva and the center of the odor source. The navigation of the *Or42a*-functional larvae yielded an average distance to the source significantly lower than that observed in presence of the solvent alone (Figure 2Diii). This result is consistent with the behavior of wild-type larvae in response to attractive odors (33). It establishes that PiVR can automatically detect and accurately track animal in real time.

Next, we tested the ability of PiVR to create virtual olfactory realities by optogenetically stimulating the larval olfactory system. In past work, robust chemotaxis was elicited in light gradients by expressing the blue-light-gated ion channel, channelrodopsin, in the *Or42a-* expressing OSN of blind larvae (17). In these experiments, the light stimulus was delivered as a point of LED light kept focused on the larva. Using a closed-loop paradigm, the intensity of the LED light was updated at a rate of 30 Hz based on the position of the larva’s head mapped onto a landscapes predefined by the user (17). PiVR was built on the same principle with the following modifications: (i) the spatial resolution of PiVR was reduced due to the fact that the field of the view of the camera captures the whole arena and not just the larva; (ii) optogenetic stimulation was achieved through homogeneous background illumination instead of a light spot that must follow the larva; and (iii) we favored the red-light-gated-ion-channel CsChrimson (22) over channelrhodopsin to minimize the innate photophobic response of larvae to the blue light range (34).

The simplified hardware design of PiVR produced accurate tracking of the head position of a larva exposed to a fictive light gradient (Figure 2B). The spatio-temporal resolution of the tracking is illustrated in Figure 2C where spikes in head speed are associated with scanning movements of the head on a timescale shorter than 500 ms (35). Head “casts” induced transient changes in light intensity *I*(*t*) (17). These changes in stimulus intensity correspond to spikes in the relative sensory experience of the larva 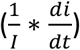 (Figure 2C, arrowheads in purple trace) (29). The tracking resolution of PiVR enabled recording periodic patterns in tail speed (Figure 2C, top green trace) that reflect consecutive cycles of acceleration/deceleration during forward peristalsis (36). *Or42a*-functional larvae displayed strong chemotaxis in response to a point-source virtual-odor gradient (Figure 2E and S3 Movie) with a level of attraction comparable to the behavior evoked by a real-odor gradient (Figure 2D). Moreover, PiVR recapitulated the meandering trajectories along the rim of a volcano-shaped virtual-odor gradient (Figure 2F), as was originally observed by Schulze, Gomez-Marin (17). Together, the results of Figure 2 and Supplementary Figures S7 validate that PiVR has the tracking accuracy and closed-loop performances necessary to elicit genuine navigation responses in virtual-odor gradients.

### Adult flies avoid activation of bitter-sensing neurons by modulating their locomotion speed

After having established the capability of PiVR to create virtual realities in *Drosophila* larvae, we sought to generalize the application of this tool to other popular model organisms. Adult *Drosophila* are covered by a thick and opaque cuticle. Consequently, activating light-gated ion channels expressed in the sensory neurons of adult flies requires higher light intensities than in semi-transparent larvae (8, 22). The background illumination system of PiVR was modified to deliver light intensities as high as 50 μW/mm^2^ to penetrate the adult-fly cuticle (37). Despite a tenfold increase in locomotion speed between adult flies and larvae (peak speed of 12 mm/s and 1.6 mm/s, respectively), PiVR accurately monitored the motion of adult fruit flies for the entire duration of 5-min trials (Figure 3A-B). We turn to gustation to test the ability of PiVR to evoke orientation behavior in adult flies stimulated by virtual chemical gradients.

*Drosophila* demonstrates innate strong aversion to bitter taste (38). This behavior is mediated by a set of sensory neurons expressing the gustatory receptor *Gr66a* in the labial palps (38). Optogenetic activation of the *Gr66a*-expressing neurons alone is sufficient to elicit aversive responses (37). Using the closed-loop tracking capabilities of PiVR (Figure 3A), we examined taste-driven responses of flies expressing the red-light-gated ion channel CsChrimson in their *Gr66a*-expressing neurons. As we reasoned that navigation in response to taste might be less directed than navigation in response to airborne odors, we presented flies with a twodimensional landscape emulating a checkerboard (Figure 3B). In this virtual checkerboard, quadrants were associated with either virtual bitter taste (light “on”) or no taste (light “off”). Flies adapted their motion to avoid squares paired with virtual bitter taste (Figure 3B-D and S5 Movies). This result generalized the field of application of PiVR to fast-moving small animals such as walking adult flies.

To determine how flies actively avoid being exposed to bitter-tasting squares, we interrogated the spatial trajectories recorded by PiVR (Figure 3B). By correlating the stimulus dynamics with locomotion, we found that flies modulate their locomotion speed in response to excitation of their bitter-tasting neurons. When flies were located in a lit square eliciting virtual bitter taste, they moved significantly faster than when located in a dark square with no bitter taste. When flies encountered sensory relief in a dark square, they frequently stopped (Figure 3E-F). In summary, our results establish that PiVR is suitable to track and immerse adult flies in virtual sensory realities. Moreover, computational quantification of behavioral data produced by PiVR suggests that flies avoid bitter tastes by modulating their locomotion speed — a strategy called *orthokinesis* (39, 40).

### Zebrafish larvae adapt their turn amplitude to stay close to a virtual-light source

Due to its transparency and amenability to molecular genetics, the zebrafish *D. rerio* has emerged as a popular vertebrate model system to study how sensory representations and sensorimotor transformations arise from the activity in neural ensembles (2, 41). Here, we show that PiVR is a powerful tool to study the organization of orientation behavior of zebrafish larvae immersed in a virtual 2D light gradient (Figure 4A-B).

Zebrafish demonstrate innate positive phototaxis to white light (42). Already at the larval stage, individuals are capable of staying confined to virtual disks of white light (43). Using PiVR, we examined whether 5-day-post-fertilization (dpf) zebrafish larvae have the navigational capabilities to stay near a virtual white light source (Figure 4B and S6 Movie). We observed that larvae were strongly attracted by the center of the virtual white light source and that they kept returning to this position for the entire duration of the trial (Figure 4D-E and Supplementary Figure S8A).

The elementary motor patterns (actions) underlying the behavioral repertoire of zebrafish larvae can be decomposed into stops, slow and rapid swims (44). By analyzing the time series of the centroid speed recorded by PiVR, we observed periodic increases in swim speed (Figure 4C). These episodes correspond to swim bursts or ‘scoots’ (45). To examine the orientation strategy used by zebrafish larvae, we discretized trajectories into scoots. As described in previous work (44), each scoot was found to be approximately straight (Figure 4B, green segments comprised by the circles). At low light intensities, significant reorientation occurred between consecutive scoots compared to the controls (Figure 4G).

Given that the virtual landscape produced by PiVR resulted from a temporal update of the intensity of isotropic stimulation light, we can rule out the detection of binocular differences of the stimulus (46). What visual features direct the increase in turning at low light intensity? To address this question, we analyzed the turn angle as a function of the light intensity conditioned by the orientation of the scoots. When zebrafish larvae moved up-gradient (positive changes in light intensity), the turn angle was not modulated by the absolute intensity of the stimulus (Figure 4H, purple). By contrast, the turn rate increased for scoots oriented down-gradient (negative changes in light intensity) at low stimulus intensity (Figure 4H, green). Therefore, we conclude that the rate of turning of zebrafish larvae is determined by a combination of the absolute light intensity and the sign of changes in stimulus intensity.

Given the ability of zebrafish larvae to efficiently return to the peak of a light gradient (Figure 4D), we asked whether zebrafish larvae can also bias their turns towards the light source. To this end, we defined the turn index (β) to quantify the percentage of turn directed toward the light gradient (Figure 4I and Methods). This metric led us to conclude that zebrafish larvae do not bias their turns towards the source more often than the control (Figure 4J). In summary, our results demonstrate that PiVR can track zebrafish larvae at sufficiently high spatio-temporal resolution to identify individual scoot events. We show that zebrafish achieve positive phototaxis by increasing the amplitude of their turns when they are moving down-gradient and experiencing low light intensities. This new aspect of the sensorimotor strategy controlling zebrafish phototaxis has also been reported in an independent, recent study featuring closed-loop light stimulations in a virtual-reality paradigm (47).

### Probing the limits of orientation behavior in the larva with dynamic virtual realities

In the wild, animals rarely encounter static odor gradients. Due to the turbulent nature of the atmosphere, natural olfactory scenes experienced by flying insects are discontinuous (48, 49). Airborne odors are spatially distributed as discrete filaments or plumes. Until recently, our field was lacking adequate technology to study how *Drosophila* converts the detection of turbulent odor plumes into coherent orientation responses. Using smoke to visualize the diffusion of small particles in air, (50) pioneered a correlative study between the perception of turbulent odor plumes and control of orientation responses in adult fruit flies. Using the closed-loop tracking and optogenetic stimulation capabilities of PiVR, this work can now be generalized to other experimental conditions.

In the present work, we asked whether *Drosophila* larvae are capable of processing discontinuous olfactory stimuli. To this end, we subjected *Or42a*>CsChrimson larvae to a reconstruction of the turbulent plumes emitted by a real point source in which adult flies were tested (50). In the experiments conducted with PiVR (Figure 5), the use of virtual realities had the advantage of guaranteeing the same stimulus conditions across trials —a feat impossible to achieve with real odor plumes. In addition, the virtual-reality technology permitted modifying specific aspects of the dynamic olfactory environment such as the replay speed of the plumes. We compared the navigational performances of *Or42a*>CsChrimson larvae in a static landscape corresponding to a “frozen” snapshot of the odor plume (S7 Movie) with the behavior observed in a *slow* odor plume structure corresponding to a 1/15^th^ of the speed of the real odor plume (S8 Movie) as well as in a fast odor plume replayed at its original speed (S9 Movie). Our results indicate that larvae are capable of orienting in slow-changing but not in fast-changing landscapes (Figure 5).

**Figure 5:**
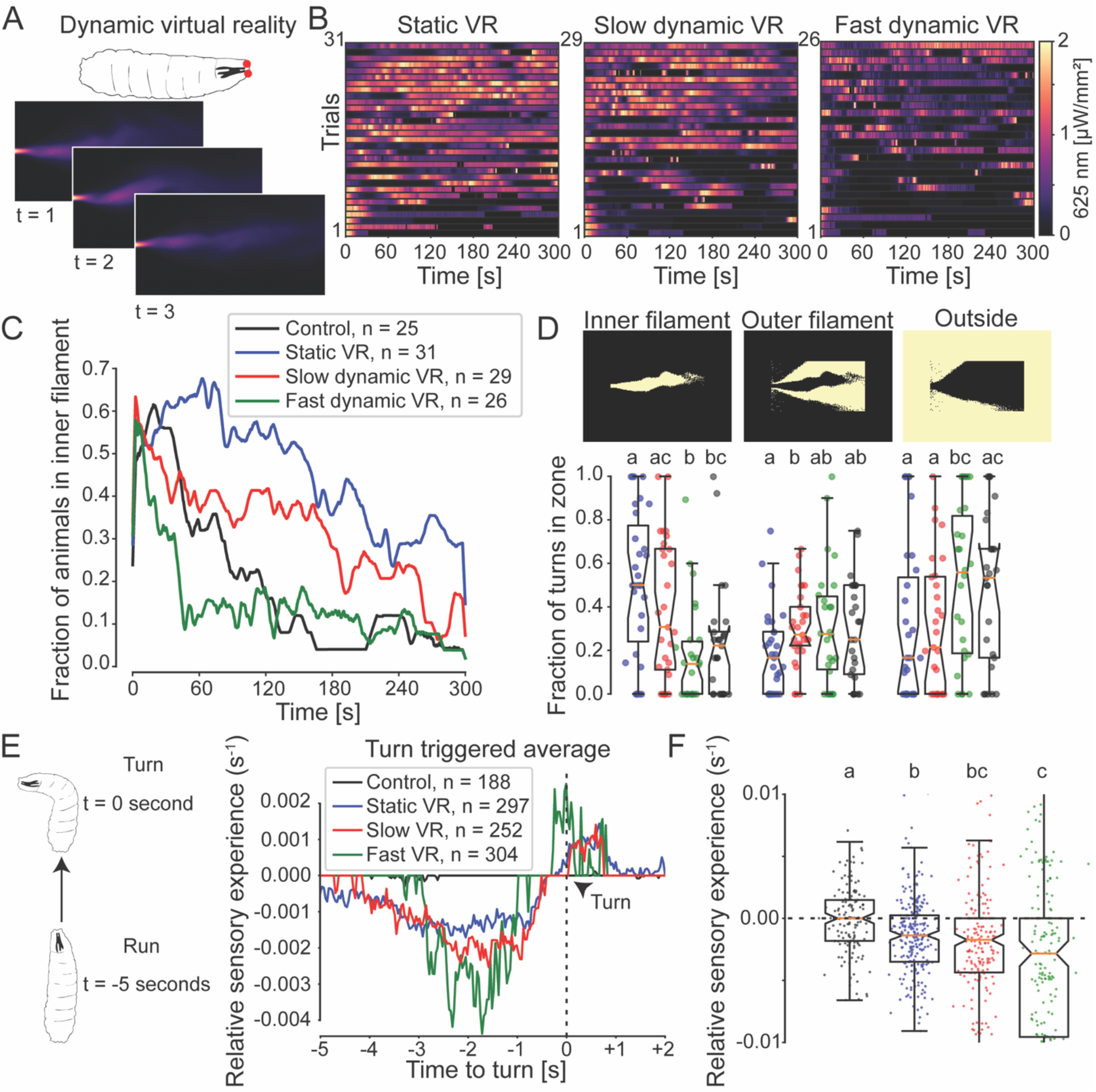
Probing the limits of orientation behavior with dynamic virtual realities. **(A)** *Drosophila* larvae expressing CsChrimson in the *Or42a* olfactory sensory neuron **(top)** were presented with a dynamic virtual-odor plume **(bottom). (B)** Ethograms of experienced light intensities for animals presented with a *Static*, a *Slow* (1/15^th^ of real speed) or *Fast* (real speed) virtual-odor plumes. Each row of the diagram represents a different trial. **(C)** Number of animals per condition in the *inner filament* (see below) over time. The *Control* genotype was presented with the *Static* virtual-odor plume. **(D) (Top)** Illustration of the *inner filament*, the *outer filament* and *outside*. The filament regions correspond to 10-100%, 2-10% and 0-2% of the maximum intensity, respectively. **(Bottom)** Fraction of turns performed in the different areas of the virtual-odor plume (Dunn’s multiple comparisons test, different letters indicate at least p < 0.05). **(E)** The relative sensory experience 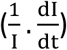 is calculated for each turn. The median of the relative sensory experience is shown for each experimental condition. **(F)** Sensory experience at 2 seconds before the turn (Dunn’s multiple comparisons test, different letters indicate cases where p < 0.01). Statistical procedures are detailed in the Methodology section. Statistical significances are indicated with lowercase letters.

To quantify the ability of larvae to ascend a turbulent olfactory landscape, we divided the plume structure into three regions: (i) the *inner filament* enclosing the medium to high intensities of the virtual-odor gradient; (ii) the *outer filament* enclosing the lower stimulus intensities and (iii) the region *outside* the plume where the virtual-odor is absent (Figure 5D, top panels). In the *static* (frozen) condition, we found that *Drosophila* larvae stayed in the inner filament region compared to genetic controls (Figure 5C, Figure 5D and Supplementary Figure S9A). Also larvae experiencing a *slow* odor plume structure (red distribution) spent more time in the inner filament compared to controls. In the *fast* odor plume structure (green distribution), larvae were unable to remain in the inner filament region longer than the controls (Figure 5C, Figure 5D and Supplementary Figure S9A). As larvae are still able to stop correctly when moving downgradient (Figure 5E-F), we conclude that larvae must have lost their ability to bias turns toward the gradient.

To summarize, Figure 5 provides a proof-of-concept that PiVR can be used to replay naturalistic dynamic olfactory landscapes while gaining control over different parameters of the stimulus. We find that larvae expressing the optogenetic tool CsChrimson in a single olfactory sensory neuron are capable of integrating turbulent olfactory signals to release turning maneuvers during down-gradient runs and to orient turns toward the local gradient. As the replay speed of the plume increases, the time during which the geometry of the local odor gradient remains correlated decreases. When the speed of the plume diffusion exceeds that of the turn dynamics, chemotaxis becomes largely compromised.

## Discussion

Anyone seeking to crack the neural logic underlying a navigation process —whether it is the response of a cell to a morphogen gradient or the flight of a seabird guided by the Earth’s magnetic field— will face the need to characterize the basic orientation strategy before speculating about its molecular and cellular implementation. PiVR is a versatile closed-loop experimental platform devised to assist the study of orientation behavior and neural-circuit functions. PiVR was developed to create virtual sensory realities by tracking the motion of unconstrained small animals subjected to a predefined model of the sensory environment (Figure 1). By capitalizing on the latest development of molecular genetics and bioengineering (3), virtual sensory stimuli are produced by expressing optogenetic tools in targeted neurons of the peripheral nervous system an animal. In the present study, we illustrate the use of PiVR by immersing *Drosophila* at the larval and adult stages in static (Figures 2 and 3) and dynamic (Figure 5) virtual chemosensory gradients and by testing the behavior of zebrafish larvae in a virtual-light gradient (Figure 4).

### An experimental platform that is both affordable and customizable

Prior to the commercialization of consumer-oriented 3D printers and cheap microcomputers such as Raspberry Pi, virtual-reality paradigms necessitated using custom setups that cost several thousand or even a hundred thousand dollars. The Raspberry <u>Pi</u> <u>V</u>irtual <u>R</u>eality system (PiVR) is an affordable (< $500) alternative that enables laboratories without advanced technical expertise to conduct sophisticated and high-throughput virtual-reality experiments. The creation of realistic immersive virtual realities critically depends on two parameters: the update frequency and the update latency of the closed-loop system. The maximal update frequency corresponds to the maximally sustained frame rate that the system can support. The update latency is the latency between an action of the tested subject and the implementation of a change in the virtual-reality environment. In normal working conditions, the update frequency of PiVR is 50 Hz while the update latency is 10-20ms (Supplementary Figure S2). These characteristics are similar to those routinely used to test optomotor responses with visual display streaming during walking behavior in insects (typically 20 Hz, 20), thereby ensuring the system’s suitability for a wide range of applications.

Different optogenetic tools require excitation at different wavelengths ranging from blue to deep red (22, 51, 52). The modularity of PiVR enables the experimenter to customize the illumination system to any wavelength range. Additionally, animals demand distinct levels of light intensities to ensure adequate light penetration in transparent and opaque tissues. While 5 μW/mm^2^ of red light had been used to activate neurons located in the leg segments of adult flies with CsChrimson (37), 1 μW/mm^2^ is enough to activate olfactory sensory neurons of the semi-transparent *Drosophila* larva (Figure 2). In its standard version, PiVR can emit red-light intensities as high as 2 μW/mm^2^ and white light intensities up to 6000 Lux —a range sufficient for most applications in transparent animals (Figures 2, 4 and 5). For animals with an opaque cuticle, we devised a higher power version of the backlight illumination system that delivers intensities up to 22 μW/mm^2^ (525 nm) and 50 μW/mm^2^ (625 nm) (Supplementary Figure S1D and Figure 3). Given that an illumination of 50 μW/mm^2^ is approaching the LED eye safety limits (International Electrotechnical Commission: 62471), it is unlikely that experimenters will want to exceed this range for common applications in the lab.

### Defining the nature of taste-driven responses in adult flies

Adult flies can walk approximately one order of magnitude faster than larvae crawl (12 mm/s and 1.6 mm/s, respectively). The detection of bitter taste elicits strong aversive spatial responses in adult flies (37). PiVR can be used to present flies with virtual realities in which space is organized according to a checkerboard where quadrants elicit or not virtual bitter tastes (Figure 3B). To this end, we expressed CsChrimson in the *Gr66a* bitter-taste-sensing neurons of the labial palps and forelegs. Consistent with a previous report (37), we found that flies modulate their movements to avoid staying exposed to virtual bitter taste in the illuminated squares. To define the orientation strategy underlying this aversive response, it is important to correlate stimulus input with behavioral output (53). We conducted such a correlative analysis on the set of trajectories recorded by PiVR for flies immersed in the virtual bitter-taste checkerboard (Figure 3D-F).

Historically, sensory-driven orientation strategies have been classified into two groups: (i) *kinesis*, which consists of undirected turns, and (ii) *taxis*, which consists of directed turns (39, 40). In the case of a kinesis, the directionality of the behavior results from an integration of sensory input to modulate the probability of releasing undirected turns (*klinokinesis*) and/or to modulate the speed of locomotion (*orthokinesis*). An example of kinesis is the response of lamprey larvae to light, which above a stimulus threshold swim in a random direction while they stay idle otherwise. Through this orthokinesis strategy, lamprey larvae accumulate in dark regions of the environment (54). Bacteria ascend attractive chemical gradients by exploiting a klinokinesis principle; when the cell moves up-gradient (positive changes in concentration), the probability of implementing an undirected turn is reduced compared to the rate observed during down-gradient or in the absence of stimulus changes (55).

When immersed in a virtual bitter-taste checkerboard, adult flies readily located the “safe” squares devoid of virtual bitter taste. By analyzing the trajectories recorded by PiVR (Figure 3B and 3D), we discovered that the optogenetic stimulation of the *Gr66a* bitter-taste sensing neurons make flies move faster (Figure 3E-F). As flies enter a safe dark quadrant, they slow down and even stop (Figure 3B). This speed modulation is the hallmark of an orthokinesis strategy. The contribution of other orientation mechanisms that integrates spatial information is left to be examined in future work. For instance, it is possible that flies implement directed turns upon their entry in a bitter-tasting quadrant (Figure 3B and S5 Movie).

Remarkably, the optogenetic stimulation of the *Gr66a* bitter-taste-sensing neurons produces a reduction in locomotion speed at the larval stage. The perception of aversive taste during larval foraging triggers stopping and reorientation (56). Thus, it is reasonable to speculate about the existence of stage-specific differences in the sensorimotor responses triggered by bitter tastes at the larval and adult stages. While the *Gr66a*-expressing gustatory neurons of the larva are likely to be presynaptic to a descending pathway commanding stopping (33), these gustatory neurons might also innervate different descending pathways controlling locomotion speed.

### Exploring the ability of zebrafish to orient with minimal graded visual inputs

Zebrafish larvae move approximately as fast as adult flies. However, their locomotor behavior is more discontinuous as it consists of bouts of locomotion with large accelerations followed by long deceleration phases (Figure 4). While these conditions make tracking more challenging, PiVR successfully monitored zebrafish larvae for several minutes (Figure 4 and S6 Movie). The zebrafish larva strongly relies on its visual system to forage, thereby making it a premier model system to study optomotor responses (57). Accordingly, we studied the orientation strategy of the zebrafish larva in virtual-light gradients.

Zebrafish are innately attracted by (real) sources of white light (42). Positive phototaxis results from an alternation between turns towards the light source and forward swims. Turns are partly directed by the detection of intensity differences measured between the left and right eyes (42, 58). To test the potential contribution of temporal changes in light intensity to the orientation process, Chen and Engert (43) presented larval zebrafish with a virtual white light disk. In spite of the fact that this landscape only featured all-or-none changes in light intensity, zebrafish larvae managed to stay inside the virtual-light disk. This spatial confinement arose from an increase in turning following the fish’s exit from the virtual-light disk (klinokinesis, 39, 40). In the present study, we used PiVR to ask whether the same mechanism can guide phototaxis in a light gradient with a Gaussian shape similar to the landscapes tested by Burgess, Schoch (42).

We presented zebrafish larvae with a virtual white light gradient. As in the work of Chen and Engert (43) and more recently Karpenko, Wolf (47), PiVR delivered a homogeneous light that precluded zebrafish from detecting binocular differences in light intensities. Reorientation in the virtual-light landscape of Figure 4 could only be guided by the detection of temporal changes in light intensity. In contrast with the all-or-none changes in light intensities produced by light disks (43), the Gaussian gradient evoked graded changes in stimulus (Figure 4C). Therefore, our results show that zebrafish larvae are capable of staying close to the source of a virtual white light source.

In a real white light gradient, fish turn towards the light source by comparing changes in light intensity experienced between their two eyes (42, 47). Due to the homogenous stimulation illumination in our experiments, fish could not use this strategy. Instead, we found that zebrafish larvae did modulate their turn angle as a function of the absolute light intensity (Figure 4G). Consistent with the results of a recent study (47), we found that the turn angle tended to increase after down-gradient scoots (Figure 4H). This observation is reminiscent of the increase in turn angle after leaving a virtual circle (43). Moreover, negative changes in light intensity did not produce an increase in turn amplitude when the absolute light intensity was sufficiently high (Figure 4H). Finally, zebrafish larvae were unable of biasing their turns toward the peak of virtual-light gradients (Figure 4J).

Together, our results highlight a new facet of the sensorimotor control of phototaxis in zebrafish. Larval zebrafish are able to stay close to a virtual-light source when the visual stimulus is reduced to an elementary optic flow perceived identically by both eyes. Under these conditions, the only sensory information available to the animal are temporal changes in light intensity due to motion in the virtual landscape. This information is sufficient for zebrafish larvae to stay near an attractive light source. The underlying reorientation strategy involves an increase in the amplitude of turns when two conditions are met: (i) the animal must detect a negative change in light intensity and (ii) the absolute light intensity must be low enough. Even in the absence of bias of individual turns toward the light gradient, the combination of the previous two conditions appear sufficient to generate positive phototaxis.

### Exploring the ability of fruit-fly larvae to orient in turbulent olfactory environments

All virtual realities discussed so far were static: their spatial structure did not change over time. The use of static gradients is convenient —if not necessary— to delineate trends in sensorimotor responses. Nevertheless, animals tend to navigate dynamic sensory environments in the wild. Insects such as the hawkmoth *Manduca sexta* (49) and the fruit fly *Drosophila melanogaster* (50, 59) orient in turbulent olfactory conditions to pinpoint the source of an attractive odor. Vertebrates can solve the same sensory challenge (60). Using PiVR, we examined how *Drosophila* larvae with a single functional olfactory sensory neuron (Figure 2Ei) respond to a naturalistic odor plume structure (50). While the chemotactic proficiency of *Drosophila* larvae has already been documented (27, 28, 32), their ability to process and orient in responses to dynamic odor plumes was unknown. By exploiting the virtual-reality possibilities offered by PiVR, we presented animals with the same dynamic plume structure at different playback speeds (Figure 5B).

As expected, larvae were able to stay within a frozen snapshot of the plume (S7 Movie). Similarly, when presented with a slowly-changing odor plume, larvae were capable of staying confined within or near the odor plume (S8 Movie). Reorientation performances strongly degraded when the replay time of the odor plume was unprocessed (Figure 5C and S9 Movie). To understand the causes of the inability of larvae to process the information associated with fast-moving plumes, we carried out a statistical analysis on the signals triggering turning maneuvers (28). For all experimental conditions, we found that the turn-triggered average of the sensory experience showed a decrease in stimulus intensity for several seconds preceding a stop (17, 56, 61) (Figure 5E). This characteristic decrease in stimulus has been shown to inhibit the activity of the olfactory sensory neurons, which in turn enhances the probability of interrupting forward peristalsis (17, 33).

Since *Drosophila* larvae were capable of releasing stops and turns during down-gradient motion in fast-changing plume structures, we reasoned that their inability to stay within the plume must be caused by a deficit in the orientation of turning maneuvers. Following a stop, larvae engage in lateral head cast to scan the local odor gradient (29). As the spatial organization of the plume changed considerably over the timescale of a head cast (~0.5 s) and a turn (~1s), larvae were unable to infer the direction of the local gradient to bias their turns toward the plume filament. As a result, they gradually drifted away from the center of the plume. In future work, it will be interesting to examine whether the combination of olfactory information with wind direction (62) improves the navigation performances of larvae in odor plumes. PiVR offers a convenient framework to study how sensorimotor mechanisms have adapted to the physical properties of natural stimuli.

### Outlook

While this study focused on sensory navigation to demonstrate the capabilities of PiVR, this tool is equally suited to crack the function of neural circuits (6) through optogenetic manipulations. To determine the connectivity and function of elements of a neural circuit, one typically performs acute excitations of specific neurons during behavior. PiVR permits the time-dependent or behavior-dependent presentation of light stimuli to produce controlled gain-of-function manipulations. The use of multiple setups in parallel is ideal to increase the throughput of behavioral screens. If the experimenter wishes to define custom stimulation rules —for instance, playing a light ramp whenever an animal stops moving— this rule can be readily included in the source code of PiVR. For pattern of light stimulation that do not depend on the behavior of an animal —for instance, stimulation with a regular series of brief light pulses—, groups of animals can be recorded at the same time. Tracking of individuals in groups can be achieved by analyzing image sequences with the basic multi-animal tracking algorithm built in PiVR or by delegating tracking to specialized algorithms such as idtracker.ai (25) (Supplementary Figure S2Bi).

Until recently, systems neuroscientists had to design and build their own setup to go about cracking the function of neural circuits or they had to adapt existing systems that were often very expensive and complex. Fortunately, our field has benefited from the publication of a series of customizable tools to design and conduct behavioral analysis. The Ethoscope is a cost-efficient solution based on the use of a Raspberry Pi computer (21), as is the case for PiVR. There is also a number of other software packages that can be used to track animals in realtime on an external computer platform. FreemoVR can produce impressive tridimensional virtual visual realities (13). MARGO is a software solution that can track a large number of animals tested in separate arenas to present individuals with closed-loop stimulation patterns (63). PiVR complements these tools by proposing a versatile and low-cost solution to carry out closed-loop tracking with low-latency performance compatible with virtual-reality experiments.

We anticipate that the performance (Supplementary Figure S1) and the resolution of PiVR (Methods) will keep improving in the future. Historically, a new and faster version of the Raspberry Pi computer has been released every two years. In the near future, the image processing time of PiVR might decrease to just a few milliseconds pushing the frequency to well above 50Hz. Following the parallel development of transgenic techniques in non-traditional genetic model systems, it should be possible to capitalize on the use of optogenetic tools in virtually any species. Although PiVR was developed for animals measuring no more than a couple of centimeters, it should be easily scalable to accommodate experiments with larger animals such as mice and rats. Together with FlyPi (64) and the Ethoscope (21), PiVR represents a low-barrier technology that should empower many labs to characterize new behavioral phenotypes and crack neural-circuit functions with minimal investment in time and research funds.

## Methods

### Hardware design

We designed all 3D printed parts using 3D Builder (Microsoft Corporation). An Ultimaker 3 (Ultimaker, Geldermalsen, Netherlands) with 0.8mm print cores was used to print all parts. We used 2.85mm PLA (B01EKFVAEU) as building material and 2.85mm PVA (HY-PVA-300-NAT) as support material. The STL files were converted to gCode using Ultimaker’s Cura software (https://ultimaker.com/en/products/ultimaker-cura-software). The printed circuit boards (PCBs) were designed using Fritzing (http://fritzing.org/home/) and printed by AISLER B.V. (Lemiers, Netherlands).

### Hardware parts

Raspberry Pi components were bought from Newark element 14 (Chicago, US) and Adafruit Industries (New York, US). The 850nm longpass filter was bought from Edmond Optics (Barrington, US). Other electronics components were obtained from Mouser Electronics (Mansfield, US), Digi-Key Electronics (Thief River Falls, US) and Amazon (Seattle, US). Hardware was obtained from McMaster (Elmhurst, US). A complete bill of materials is available in Supplementary Table S1. Updates will be available on www.PiVR.org and https://gitlab.com/LouisLab/PiVR

### Building and using PiVR

Detailed instructions on how to build a PiVR and how to use it can be found in Supplementary S_HTML1. Any updates will be available on www.PiVR.org.

### Data Analysis and Statistics

All behavioral data shown in the main and Supplementary Figures have been deposited as a data package on Dryad (*link to Dryad repository upon publication of the peer-reviewed paper)*. Data analysis was performed using custom written analysis codes which are bundled with the data package. The analysis codes are additionally available from https://gitlab.com/louislab/pivr_publication (*to be made accessible upon publication of the peer-reviewed paper)*.

### Latency measurements of PiVR

To estimate the time it takes between an animal performing an action and PiVR presenting the appropriate light stimulus, the following elements were taken into account: (I) image acquisition time, (II) image processing and VR calculations latency and (III) software to hardware latency. To measure image acquisition time (Supplementary Figure S1b), the camera was allowed to set optimal exposure at each light intensity before measuring the shutter speed. To measure image procession and VR calculations latency **(Cii)** time was measured using the non-real-time python time.time() function which we estimate has a uncertainty in the order of a few microseconds. **(Ciii)** To confirm these measurements, we also recorded the timestamps given by the real-time graphical processing unit (GPU). Combined, these two plots show that while image processing time has a median around 8-10ms, there are a few frames where the image processing time takes longer than 20ms which, at high framerates, leads to frames being skipped (arrows in **Cii** and **Ciii**). Finally, to measure the software to hardware latency we measured how long it takes to turn the general purpose input output (GPIO) pins on and off during an experiment **(Dii)**. The pins are connected to a transistor with rise and fall times in the order of μs (https://eu.mouser.com/datasheet/2/308/FQP30N06L-1306227.pdf). LEDs usually have a latency in order of 100s of ns. The total latency between an animal behaving and the appropriate stimulus being presented is therefore in the order of 10-15ms.

### Video recording performance of PiVR

When PiVR is used as a video recorder, the resolution (px) limits the maximal framerate (fps): At 640×480px the framerate can be set up to 90fps, at 1296×972 up to 42fps, at 1920×1080 up to 30fps and at 2592×1944 a framerate up to 15fps can be used. PiVR is compatible with a wide variety of M12 lenses allowing for high quality video recordings, depending on the experimental needs.

### Fruit fly larval experiments

For the experiments using fruit fly larvae (Figure 2 and Figure 5), animals were raised on standard cornmeal medium at 22°C on 12 hr-day: 12-hr-night cycle. 3rd instar larvae were placed in 15% sucrose for 20 to 120 min prior to the experiment. Experiments were conducted on 2% agarose (Genesee, 20-102). In the experiments described in Figure 2, the arena was a 100-mm diameter Petri Dish (Fisher Scientific, FB0875712). In the experiments described in Figure 5, the arena was the lid of a 96-well plate without condensation rings (Sigma-Aldrich, L3536-100EA).

For the experiments featuring a real odor gradient, isoamyl-acetate (Sigma Aldrich, 306967-100ML) was diluted in paraffin oil (Sigma Aldrich, 18512-1L) to produce a 1M solution. A single source of the odor dilution was tested in larvae with the following genotype: *w;Or42a-Gal4;UAS-Orco,Orco*^−/−^. The control consisted of solvent (paraffin oil) devoid of odor. For the virtual-reality experiments, the following genotype was used: *w;Or42a-Gal4,UAS-CsChrimson;UAS-Orco,Orco*^−/−^. In Figure 2, the same genotype was used in controls but the tests were conducted without any light stimulations. In Figure 5, the control genotype was Orco^*−/−*^ tested with the *static* odor plume. Larvae expressing CsChrimson were grown in complete darkness in 0.5M all-*trans* retinal (R2500, MilliporeSigma, MO, USA).

### Adult fruit fly experiments

Male flies with the *Gr66a*-Gal4 transgene (Bloomington stock number: 57670) (65) were crossed to virgin females carrying *20xUAS-CsChrimson-mVenus* (22) integrated into the *attP40* landing site. The flies were grown in complete darkness on standard cornmeal medium with 0.5M all-*trans* retinal at 25°C. Female flies between 1 and 7 days after eclosion were selected after putting the vial on ice for a few seconds. The experiment was conducted in a 100-mm diameter Petri Dish (Fisher Scientific, FB0875712) under white light condition.

### Zebrafish larva experiments

In the experiments shown in Figure 4, we used AB *casper* (66) as parents. Only pigmented larvae were used for the experiments. The larvae were reared at 28.5°C and a 14:10 light cycle. The experiments were run at 26°C. The arena was a 100mm Petri Dish (Fisherbrand, 08-757-12PK). All ambient light was blocked. The maximum white light intensity provided by PiVR was measured to be approximately 6800Lux (Extech Instruments Light Meter 401025).

### Data analysis and statistical procedures

All data analysis was performed using Python. The scripts used to create the plots shown in the figures (including all the data necessary to recreate the plots) can be found at (*link to Dryad repository upon publication of peer-reviewed the manuscript)*. Generally, datasets were tested for normal distribution (Lilliefors test) and for homogeneity of variance (Levene’s test) (67). Depending on the result, the parametric T-test or the non-parametric Mann-Whitney U ranksum test was used. To compare multiple groups, either Bonferroni correction was applied after comparing multiple groups or Dunn’s test was applied (67). Below, information about the analysis and the applied statistical tests throughout the manuscript are separately addressed for each figure. To estimate peak movement speed of different animals, the median of the 90^th^ percentile of maximum speed per experiment was calculated.

**Data analysis of Figure 2:** To calculate movement speed, the X and Y coordinates were first filtered using a triangular rolling filter with a window size equal to the frame rate (30 fps) divided by the high bound on the speed of the animal (1 mm/s) times the pixel-per-mm value of the experiment. Depending on the exact distance between camera and the arena (pixel-per-mm value), the window size of the filter was typically 0.3 s. Speed was calculated using Euclidian distance. The time series of the speed was smoothened using a triangular rolling filter with a window size of 1 s. To calculate the sensory experience, the stimulus intensity time course was filtered using a boxcar rolling filter with window size 1 s (Figure 2C). To calculate the distance to source for the isoamyl acetate gradient, the source location was manually defined using the PiVR software. In the Gaussian virtual-odor gradient, the coordinates with the maximum intensity value was defined as the source. In the volcano-shaped virtual gradient, the circle with the highest values was defined as the nearest source to the animal (Figure 2F). At 4 minutes into the experiment the distance to source between the experimental and control condition was compared. As the data was not normally distributed (Lilliefors test), Mann-Whitney U test was used (Supplementary Figure S7A-C).

**Data analysis of Figure 3:** Each experiment lasted 5 minutes. The preference index for each animal was calculated by subtracting the time spent by an animal in the squares without light from the time spent in the squares with light (squares eliciting virtual bitter taste). This subtraction was then divided by the total experimental time (Figure 3C). Mann-Whitney U test was used to compare preference between genotypes as the distribution of the preference indices was not normally distributed (Lilliefors test). Speed was calculated as described in Figure 2: First, the X and Y coordinates were smoothened using a triangular rolling filter with a window size equal to the frame rate (30) divided by the approximate walking speed of flies (~5 mm/s) times the pixel-per-mm ratio of the experiment, which was usually 0.06-0.09 s. Locomotion speed was calculated by using the Euclidian distance. The time series of the speed itself was filtered using a triangular rolling filter with a window size equal to the frame rate (30 fps). To test for statistical difference in speed between genotypes and the light on and off condition, Dunn’s Multiple Comparison Test was implemented through the scikit-posthocs library of Python.

**Data analysis of Figure 4:** Each experiment lasted 5 minutes. Each trial started with the animal facing the virtual-light source. To focus the analysis on animals that demonstrated significant movement during the experiment, the final dataset was based on trials fulfilling the following criteria: (1) an animal had to move at least 20 mm over the course of the experiment; (2) only movements above 1 mm/frame were recorded (due to camera/detection noise the centroid in resting animals can move); (3) the animal must have been tracked for at least 3 out of 5 minutes.

Locomotion speed was calculated by first smoothening the X and Y centroid coordinates with a half-triangular rolling filter with the window size of 1 s. The speed was calculated using Euclidian distance and filtered again using a 0.3 s triangular rolling filter. Scoots were identified from the time course of the movement speed by using the “find_peaks” function of the scipy library (68) with the following parameters: a minimum speed of 2.5 mm/s and a minimum time between two consecutive peaks (scoots) of 5 frames (0.165 s) (Figure 4C). The same filter was applied (half-triangular rolling filter with window size of 1 s) to calculate the distance to source. Distance to the maximum value of the virtual-reality landscape was calculated. To calculate the distance to source of the controls, the same virtual-reality was assigned relative to the animal starting position and the distance to this simulated landscape was calculated (Figure 4E). In Supplementary Figure S8A, the distance to source was compared across experimental conditions at 4 min into the experiment. We used Mann-Whitney U test to compare the distance to source as the variance between samples could not be assumed to be equal (Levene’s test) (Supplementary Figure S8A).

Scoots (see above) were then used to discretize the trajectory. The reorientation angle θ was calculated for each pair of consecutive scoots by comparing the animal’s coordinates 0.66 s (approximate duration of a scoot) before and after the local peak in locomotor speed. To filter out occasional (<10%) misdetection of the animal’s reflection once it was located near the wall of the petri dish, we only considered scoots with a reorientation angle θ smaller than 135° (Figure 4F). The light intensity experienced by the animal was used to bin the reorientation angles into 4 groups shown in Figure 4G-H. To compare the control vs. experimental condition of the four light intensity bins, Student’s t-test was used after checking for normality (Lilliefors test) and for equality of variance (Levene’s test). Bonferroni correction was used to correct for multiple comparisons (Figure 4G).

To compare reorientation (turn) angles θ associated with positive and negative visual sensory experiences, the light intensity change experienced during the previous scoot: (ΔI) was used to split the experimental data in two groups: ΔI>0 (upgradient) and ΔI<0 (downgradient). The data was binned according to stimulus intensity as in Figure 4G. To compare turn angle in the two groups, student’s t-test for paired samples was used after checking for normality (Lilliefors test) and for equality of variance (Levene’s test). Bonferroni correction was used to adjust for multiple comparisons (Figure 4E).

The turn index β was calculated based on the angular difference a between the heading the animal at a given scoot and the bearing with respect to the white light source. This angular difference a indicates whether the animal approaches the source (angles near 0°) or swims away from it (angles near 180°). We defined 90° as the boundary between swimming towards and swimming away from the source. The turn index β was then defined by counting the number of scoots towards the source and by subtracting to it the number of scoots away from the source normalized by the total number of scoots. A turn index was calculated for each trial. We compared each group using the Mann-Whitney U test as the data was not normally distributed (Lilliefors test). Bonferroni correction was used to adjust for multiple testing (Figure 4j).

To bin the reorientation (turn) angle θ according to distance to source, the distance was calculated for each scoot and binned in 4 groups according to Supplementary Figure S8C. We compared each group using Mann-Whitney U test as the data was not always normally distributed (Lilliefors test). Bonferroni correction was used to adjust for multiple comparisons.

**Data analysis of Figure 5:** The 2D dynamic odor plume structure (movie) was adapted from (50). Due to the limited random-access memory (RAM) available on the Rasperry Pi the data was downscaled (i) by converting the data to 8 bit and (ii) by only using the first 30 seconds of the original movie which were played in a loop. Not all trials were used in the data analysis. The inclusion criteria for individual trials were as follows: (i) At the start of the experiments the animal had to move towards the virtual source ±45° and (ii) animals had to move a minimum of 50% of the median distance travelled by all animals. The first criterion was used to filter out larvae that were not exposed to the plume at the onset of the trial. The second criterion was used to filter out a minority (<5%) of larvae that were abnormally inactive.

To score the ability of animals to stay in a virtual-odor plume, the 2D plume structure was decomposed into three regions: the *inner filament*, the *outer filament* and the area *outside* the filament (Figure 5D). The *Inner filament* was defined as the region were the light intensity was comprised between 10% and 100% of the maximum intensity. The *outer filament* was defined as the region where the light intensity was between 2% and 10% of the maximum intensity. The *outside* included every position with an intensity below 2%. As the odor plume moves, the boundary of these regions change during the experiment. As the behavioral quantification shown in Figure 5C aimed to characterize navigation in responses to the virtual odor, animals were labeled as inside the *inner filament* only if they had not been *outside* earlier during the trial. In this analysis, the behavior of the animal was ignored upon first exit of the filament.

To compare the ability of larvae to stay in the *inner filament*, Dunn’s multiple comparison test was used (Supplementary Figure S9A). To quantify the turning maneuvers, the body angle was used according to the procedure detailed in (69). Briefly, temporal changes in body angle (angular speed) were used to classify turning maneuvers: If the change in body angle was comprised between 7 and 143 °/s for a minimum of 0.33s, the behavior was classified as a turning maneuver. Dunn’s multiple comparison test was used to compare fractions of turns in different regions of the filament (Figure 5D) and median changes in relative sensory experience observe two seconds before the turn onset (Figure 5E and Supplementary Figure S9B).

## Acknowledgement

We thank Ellie Heckscher, Primoz Ravbar and Andrew Straw for comments on the manuscript. We are grateful to Ajinkya Deogade for work performed during the initial phase of the project (development of a do-it-yourself open-loop tracker based on FlyPi), for creating a preliminary draft of 3D printed parts, for measuring the white light intensities used in Figure 4, and for discussions. We thank Tanya Tabachnik for technical advice about the assay construction. We are grateful to Stella Glasauer for collecting the Kelp Fly tracked in Figure S6A, for taking the picture of the spider in Supplementary Figure 6B, for designing the PiVR logo and for comments on the manuscript. We are in debt to Tyler Sizemore for collecting the firefly and pill bug tracked in Supplementary Figure 6. We thank Igor Siwanowicz for providing the pictures of the firefly and pill bug of Supplementary Figure 6. We are grateful to Minoru Koyama and Jared Rouchard for rearing and providing the fish used in Figure 4. The development and optimization of PiVR greatly benefited from feedback received during the Drosophila Neurobiology: Genes, Circuits & Behavior Summer School in 2017, 2018 and 2019 at the Cold Spring Harbor Laboratory.

