## Supplementary information for "PiVR: an affordable and versatile closed-loop platform to study unrestrained sensorimotor behavior"

### Supplementary figures

Supplementary Figure 1: **Timing performance of PiVR**

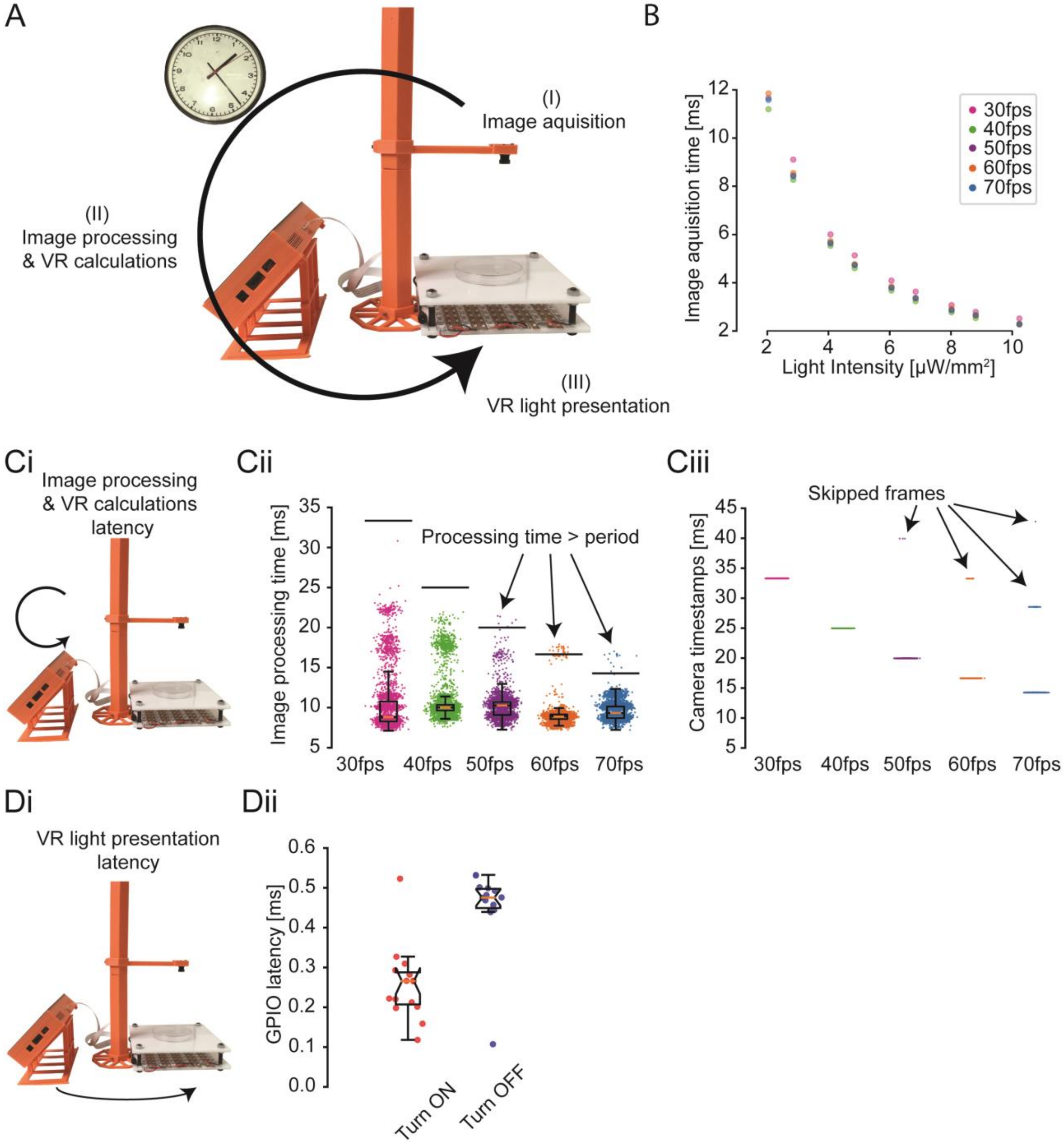

**(A)** Illustration of the three parameters measured to estimate latency. **(B)** Dependence of image acquisition time on the infrared background illumination strength. **(Ci)** image processing & VR calculations is measured first by the non-real-time python 3.5 time.time() function. At 50, 60 and 70 fps some frames (6/5598, 25/7198 and 16/8398 respectively) take longer than the period of the framerate which leads to dropped frames **(Cii)**. To confirm these measurements, we also recorded timestamps of the images assigned by the real-time clock of the graphical processing unit (GPU), as shown in panel **(Cii)**. **(D)** To estimate the software to hardware latency during a full update cycle of the tracker, we measured the time between the general purpose input/output (GPIO) pin being instructed to turn on and the GPIO pin reporting being turned on.

Supplementary Figure 2: **Modularity of PiVR (hardware)**

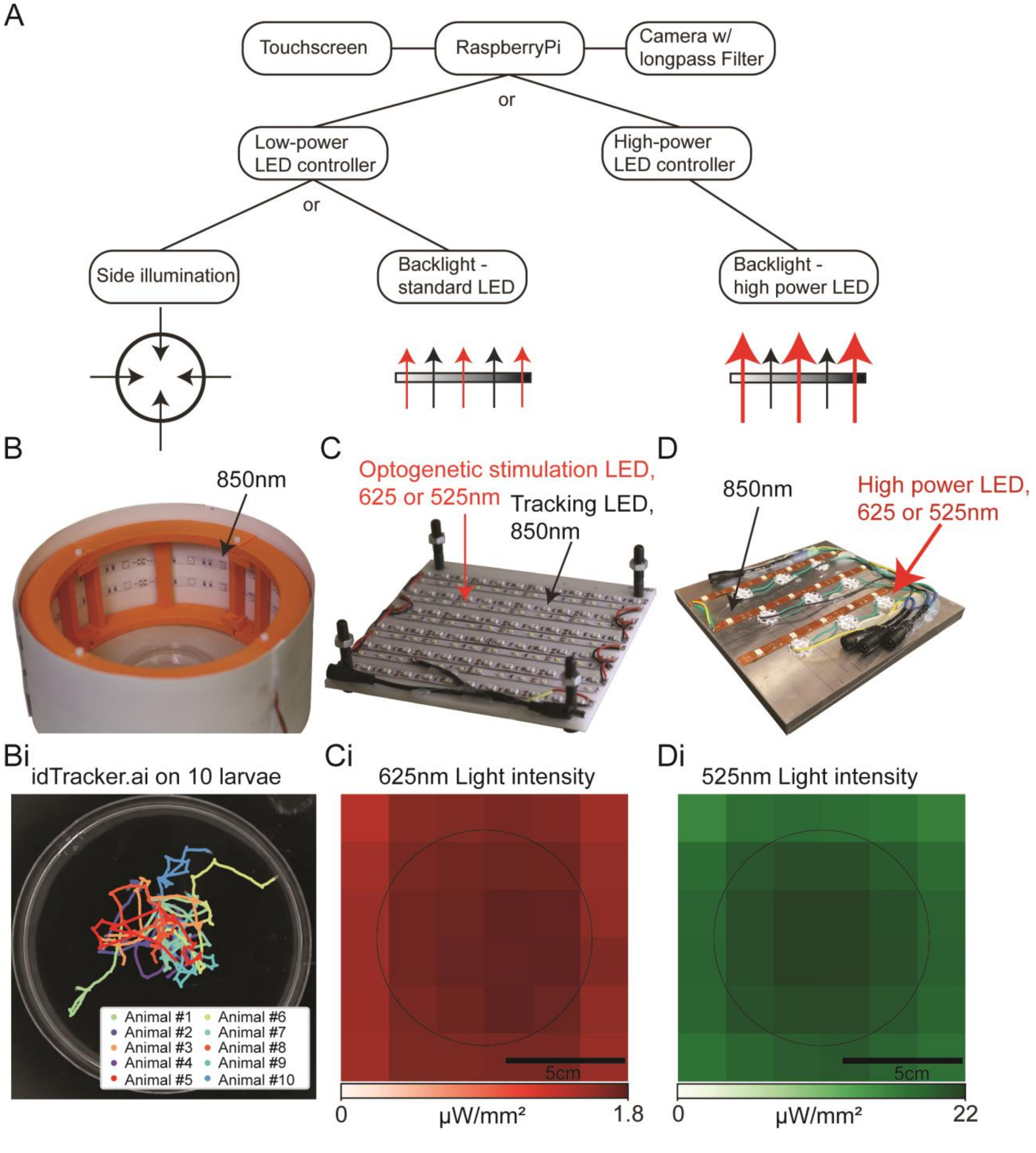


**(A)** PiVR is highly modular. The standard version (shown in Fig. 1A) can easily be adapted to allow for side (or top) illumination using the same LED controller system. If high-light intensities are needed, a high power LED controller can be used in combination with high-power LEDs. Panel **(B)** shows an example of a side illuminator. Side illumination increases contrast on the surface of the animal. **(Bi)** This side illuminator was used to collect videos with sufficient detail for idtracker.ai [1] to track 10 fruit fly larvae while retaining their identity. **(C)** The standard PiVR illuminator consists of at least two different 12V LED strips: one for background illumination (850nm) and another color (here 625nm) to stimulate the optogenetic tool of choice. **(D)** The high power stimulation arena uses a LED controller that keeps current to high power LEDs constant and can drive a maximum of 12 high power LEDs. **(Ci** and **Di)** The intensity of the stimulation light was estimated by using a Thorlabs Inc. S130VC. Each pixel is 2 $cm^2$. The black circle indicates petri dishes used as behavioral arenas.

Supplementary Figure 3: **Flow diagram of the automatic animal detection and background reconstruction**

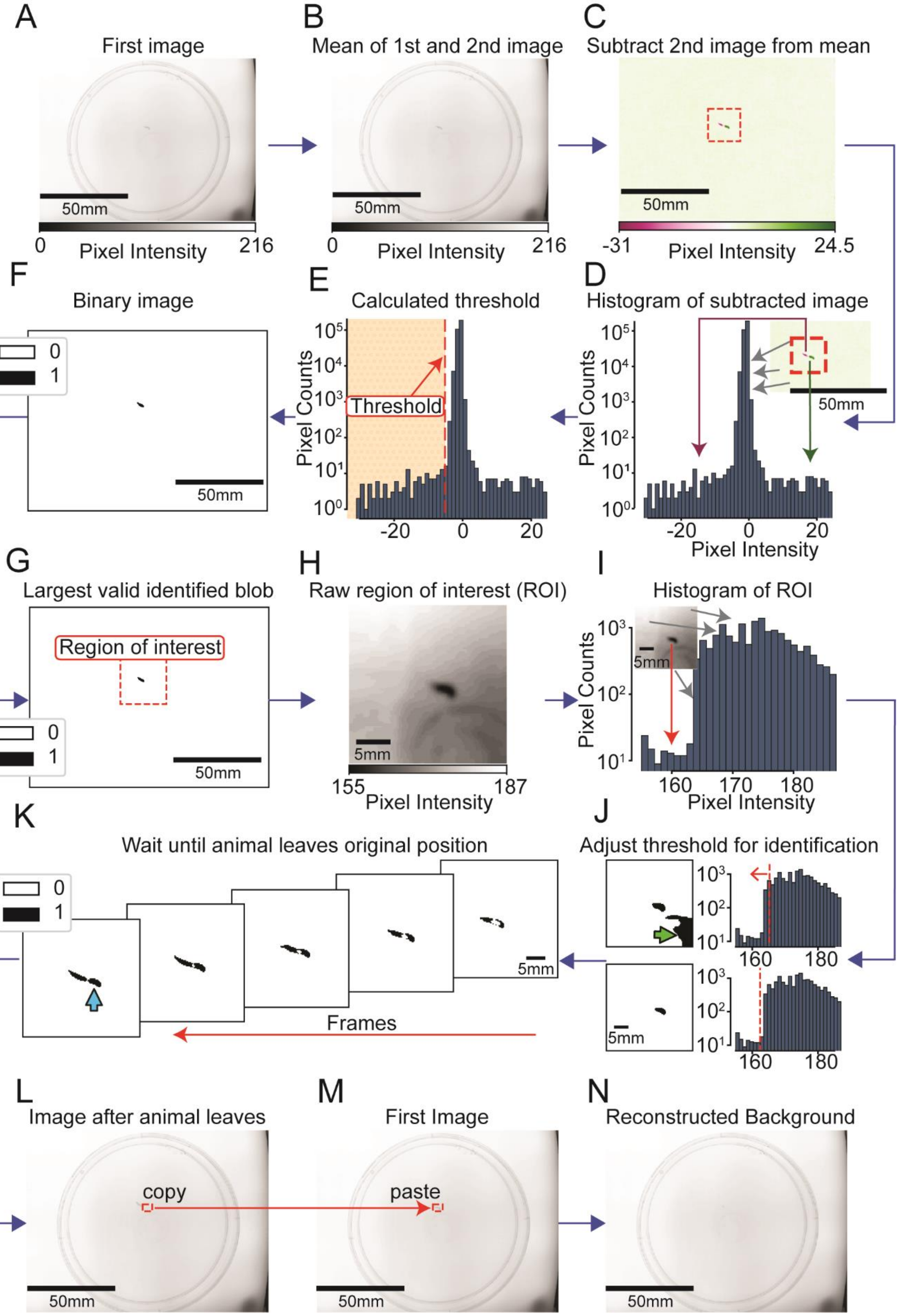

**(A)** After placing the animal and pressing ‘start tracking’ on the graphical user interface, the software will grab the first image. All images are immediately filtered using a Gaussian kernel with a sigma depending on the size of the animal (user defined) to reduce camera noise. **(B)** Then a second image is taken. The mean of the images taken so far is being calculated. **(C)** The current image (in this example the second image of the experiment) is then subtracted from the mean image shown in panel **(B)**. **(D)** The histogram of the subtracted image shows most greyscale values to be 0. For the region where the animal has moved since the first frame the pixel values are negative (magenta). For the region that the animal has left since the first frame, the pixel values are positive (green). **(E)** The threshold value is calculated based on the histogram: it is the mean of the image subtracted by 4 (optimal value defined by trial and error). **(F)** The threshold value is used to binarize the subtracted image shown in panel **(C)**. If there is no or more than one blob with a minimal area (defined by user in animal parameters file) the loop restarts at step **(B)**. **(G)** If there is exactly one blob, an area around the blob (defined by user in animal parameters file) is defined as the current region of interest. **(H)** The region of interest is the area where movement has been detected. The algorithm will now restrict the search for the animal to this region. **(I)** The histogram of this small area of the first image **(A)** shows that the few pixel defining the animal are distinct from the background. **(J)** To find the optimal local threshold for binarizing the image, the threshold is adjusted if more than one blob is detected (**top, green arrow**). As soon as only one blob with the characteristics of the animal (defined by user in animal parameters file) has been detected, the local threshold value is set and the shape of the identified animal in the first frame saved **(bottom)**. **(K)** Using the local threshold, each new image is binarized and then subtracted from the first image as shown in panel **(J, bottom)**. If the identical blob that was detected in panel **(J bottom)** is found in any of the new subtracted binary images (**cyan arrow**), the animal is considering as having left its original position and the algorithm continues. **(L)** The region occupied by the animal is then copied from the latest image and **(M)** pasted into the first image. **(N)** The resulting image does not contain the animal and will be used as the background image for the tracking algorithm (Supplementary Figure S4).

Supplementary Figure 4: **Flow diagram of the animal tracking algorithm**

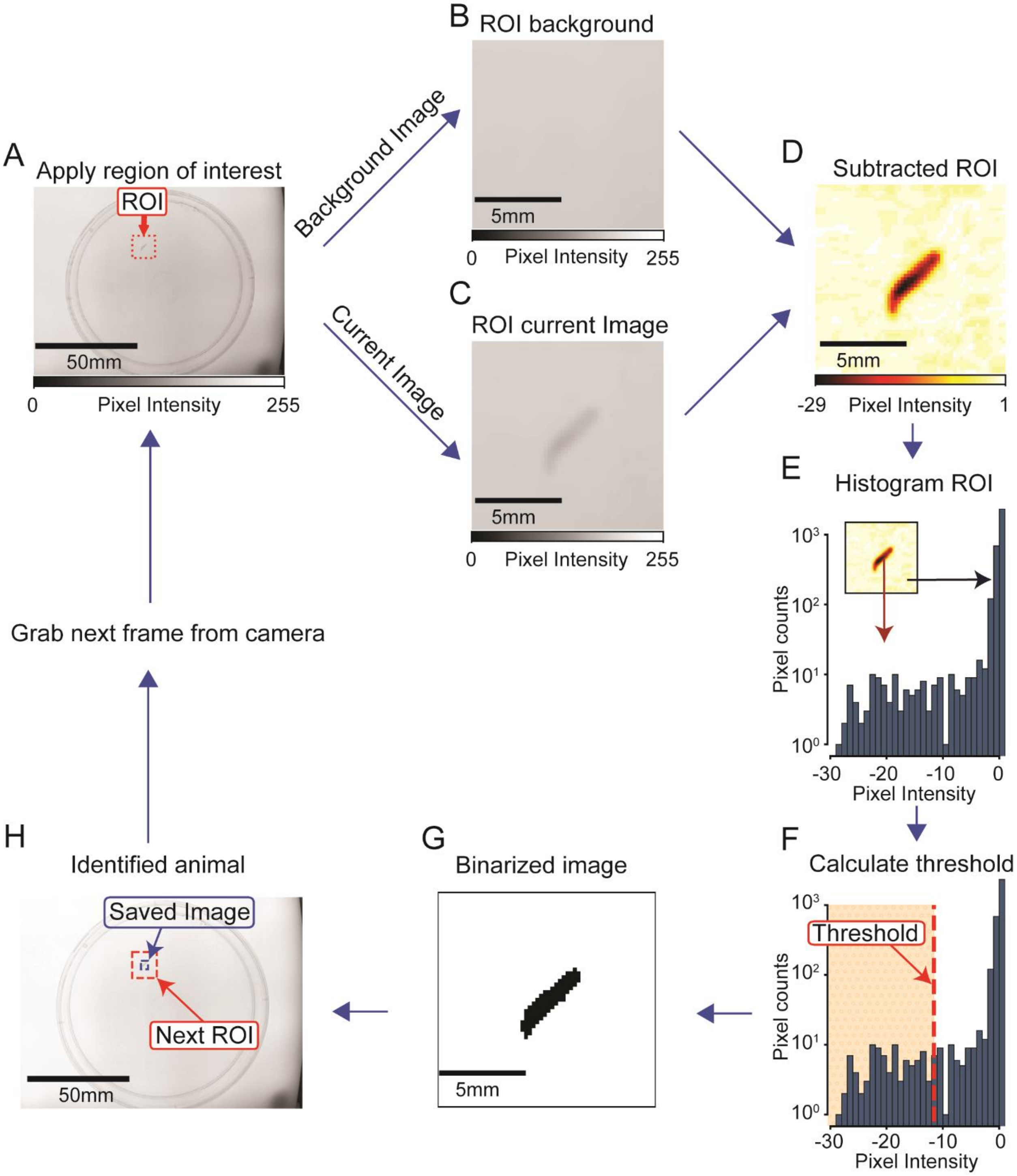


**(A)** At the start of the experiment, the region of interest (ROI) is defined during animal detection (Supplementary Figure 3G). During the experiment, the current ROI is defined using the previous frame. The ROI of the current image **(C)** is then subtracted from the ROI of the background **(B)**. The fact that the tracking algorithm only considers a subsample of the image is central to the temporal performances (short processing time) of PiVR. **(D)** In the resulting image, the animal clearly stands out relative to the background. **(E)** The histogram of the image indicates that while the background consists mostly of values around 0, the animal has pixel intensity values that are negative. **(F)** The threshold is defined as being three standard deviations away from the mean **(G)**. This threshold is used to binarize the subtracted ROI. The largest blob with animal characteristics (defined by animal parameters) is defined to be the animal. **(H)** The image of the detected animal is saved and the next ROI is designated (defined by animal parameters).

Supplementary Figure 5: **Head/Tail classification using an Hungarian algorithm**

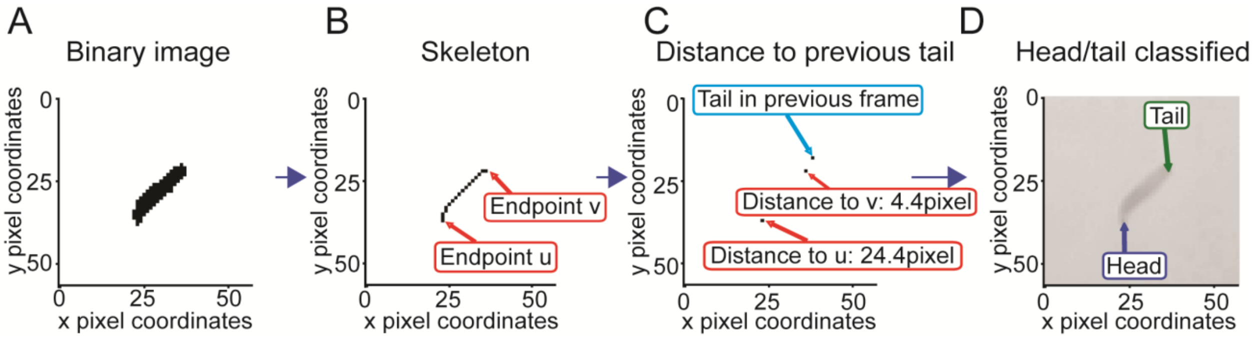


**(A)** During tracking, head tail classification starts with the binarized image (Supplementary Figure 4G). **(B)** The binary image is used to calculate the morphological skeleton which in turn is used to identify the two endpoints, one of which must be the head and the other the tail. **(C)** The Euclidian distance between the tail position in the previous frame and each of the endpoints is calculated. If the tail has not been defined in the previous frame, the centroid position is used instead. **(D)** Whichever endpoint has less distance is defined as the tail (here v). The other endpoint is defined as the head.

Supplementary Figure 6: **Capability of PiVR to track a wide variety of small animals**

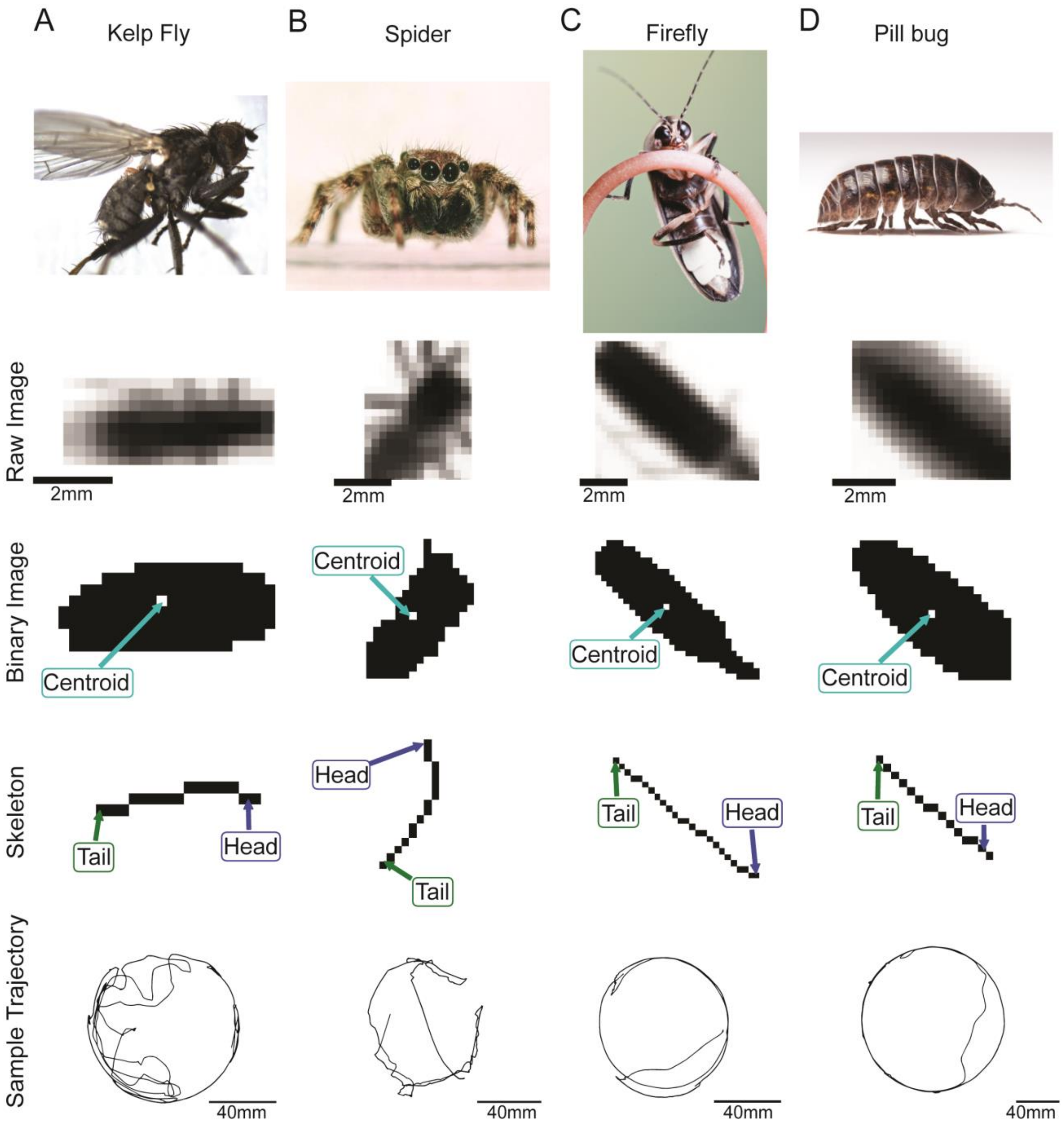


PiVR is able to detect, track and assign head and tail positions to a variety of invertebrate species with different body plans: **(A)** kelp fly, **(B)** jumping spider, **(C)** firefly and **(D)** Pill Bug.

Supplementary Figure 7: **Distance to real and virtual odor sources in larval chemotaxis to real and virtual odor gradients**

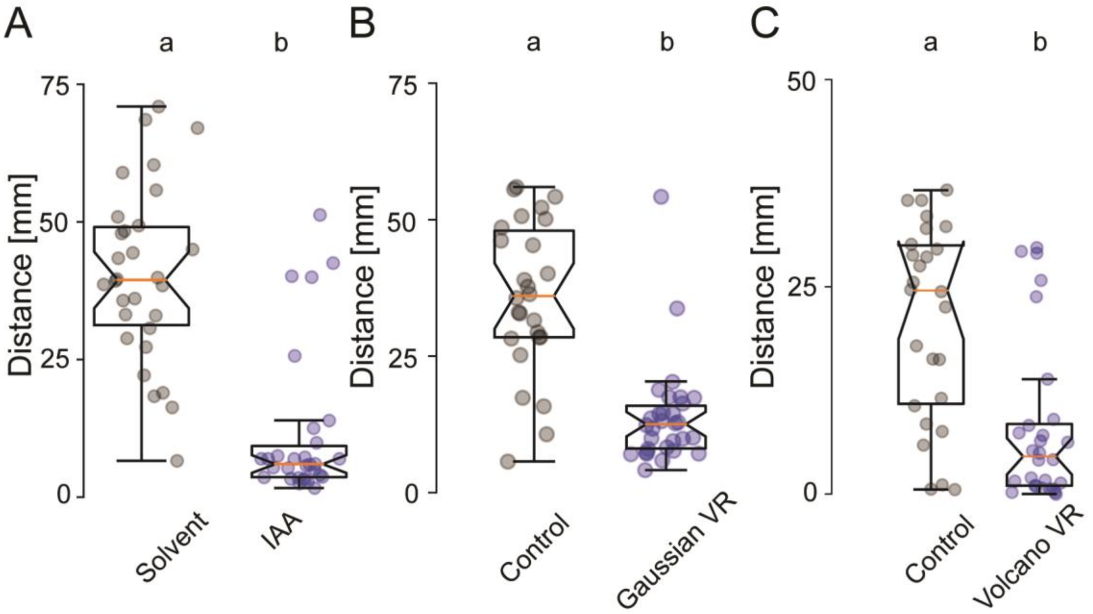


**(A)** Distance to real odor source (iso-amyl acetate, *n* = 30) and the solvent (paraffin oil, *n* = 30), **(B)** between the gaussian shaped virtual odor reality (*n* = 31) and the control (*n* = 26) and **(C)** the distance to the local maximum (rim of the volcano, *n* = 29) and the control (*n* = 26). Timepoint is 4 minutes into the experiment (Mann-Whitney U test, $p < 0.001$).

Supplementary Figure 8: **Zebrafish larvae in virtual light source**

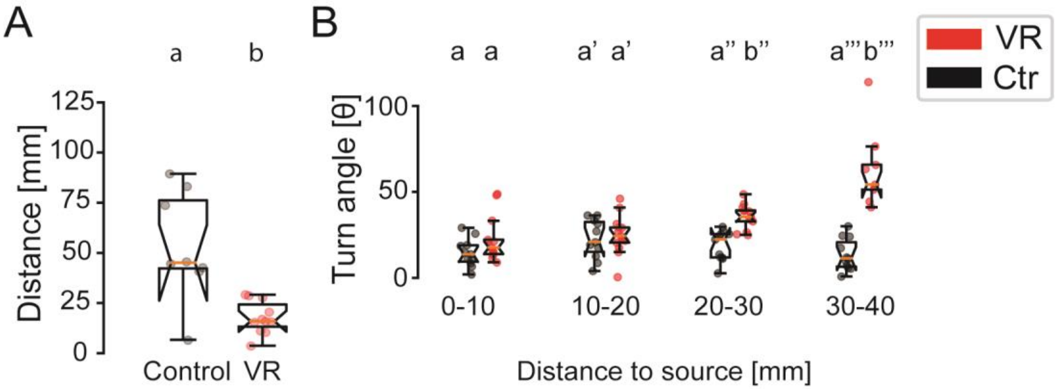


**(A)** Distance to virtual light source of control (black) and experimental condition (red) at 4 minutes into the experiment. **(B)** Relationship between turn angle θ and distance to the virtual light source (Mann-Whitney U test, different letters indicate $p < 0.01$, $n = 11$ and 13). All reported p-values are Bonferroni corrected.

Supplementary Figure 9: **Detailed analysis of inability of larvae to stay close to a fast moving virtual odor plume**

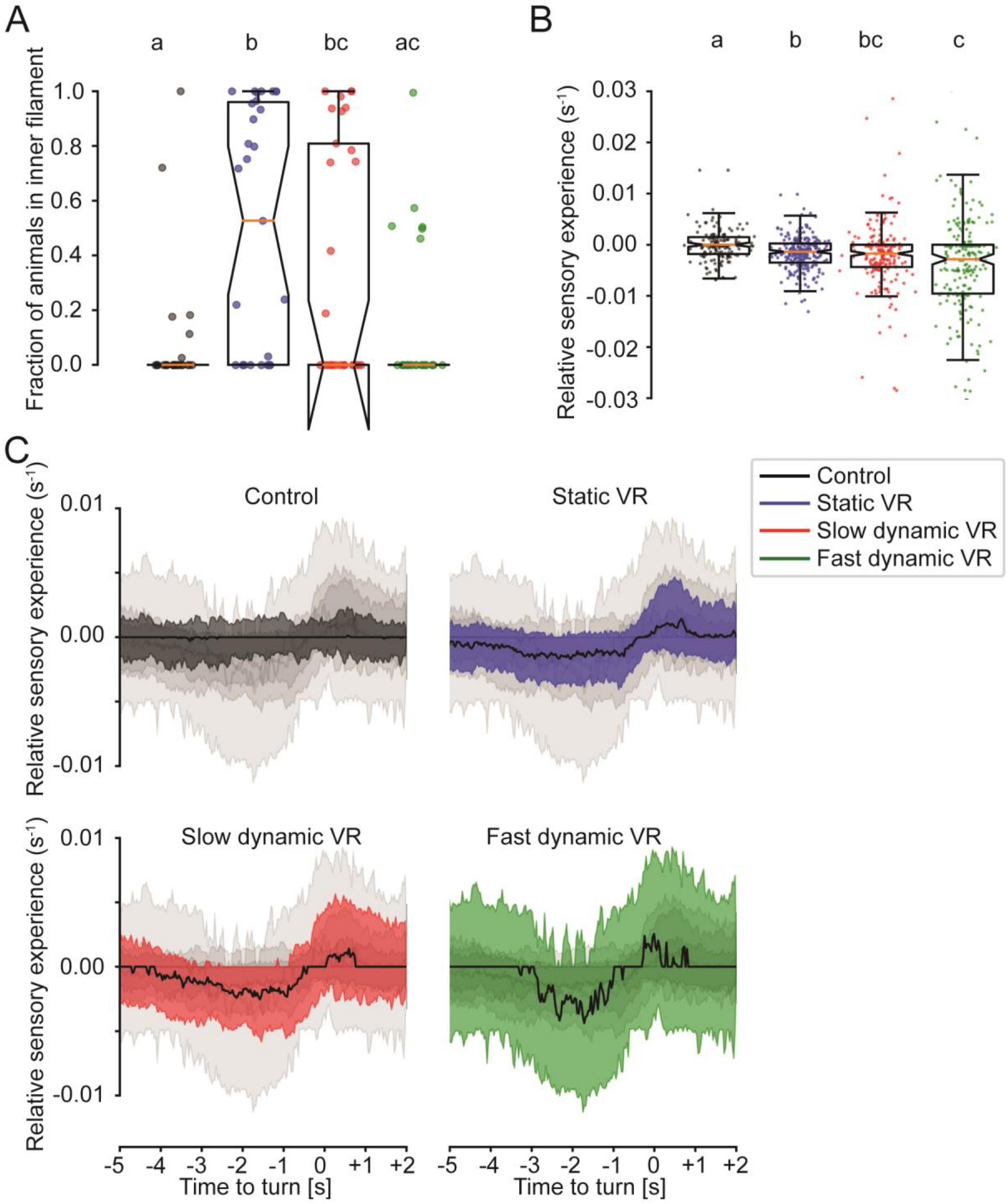


**(A)** Time courses of the fraction of animals found in the *inner filament* of the virtual odor plume between one and two minutes into the experiment (Dunn's multiple comparisons test, different letters indicate at least $p < 0.05$, $n$ = 25, 31, 29 and 26). **(B)** Relative sensory experience for each turn. Zoomed out version of Figure 5f. **(C)** Median relative sensory experience before a stop (black line) and the standard deviation for each experimental condition ($n_{control}$ = 188, $n_{static}$ = 297, $n_{slow}$ = 252 and $n_{fast}$ = 304).

### Supplementary movies

Supplementary Movie 1: Illustration of homogenous illumination creating a virtual checkerboard reality. The behavior of a freely-moving fly is shown in the petri dish (**Left**), in the virtual checkerboard recorded by PiVR (**Middle**). The corresponding time course of the homogenous illumination intensity is shown in the (**Right**) panel.

Supplementary Movie 2: Sample trajectory of a *Drosophila* larva expressing *Orco* only in the *Or42a* olfactory sensory neuron behaving in a quasi-static isoamyl-acetate gradient. The odor gradient was reconstructed for visualization and an estimation of the experienced odor intensity as described before [2] (see also Methodology section).

Supplementary Movie 3: Illustrative trajectory of a *Drosophila* larva expressing the optogenetic tool CsChrimson in the *Or42a* olfactory sensory neuron behaving in a "Gaussian" shaped virtual odor reality.

Supplementary Movie 4: Illustrative trajectory of a *Drosophila* larva expressing the optogenetic tool CsChrimson in the Or42a olfactory sensory neuron in a "volcano" shaped virtual odor reality.

Supplementary Movie 5: Illustrative trajectory of an adult *Drosophila* expressing the optogenetic tool CsChrimson in the *Gr66a* bitter sensing neurons in a "checkerboard" shaped virtual gustatory reality.

Supplementary Movie 6: Illustrative trajectory of a zebrafish larva in a virtual visual reality.

Supplementary Movie 7: Illustrative trajectory of a *Drosophila* larva expressing the optogenetic tool CsChrimson in the *Or42a* olfactory sensory neuron in a *static* virtual odor plume.

Supplementary Movie 8: Illustrative trajectory of a *Drosophila* larva expressing the optogenetic tool CsChrimson in the *Or42a* olfactory sensory neuron in a *slowly moving* virtual odor plume.

Supplementary Movie 9: Illustrative trajectory of a *Drosophila* larva expressing the optogenetic tool CsChrimson in the *Or42a* olfactory sensory neuron in a *fast moving* virtual odor plume

### Supplementary files

Supplementary Table S1: Bill of materials for the standard version of PiVR.

Supplementary HTML S1: Current version of website found at www.PiVR.org
